# Signalling-state dependent drug-tolerance in head and neck squamous cell carcinoma

**DOI:** 10.1101/2023.12.05.570063

**Authors:** Dyah W. Karjosukarso, Alice Dini, Laura J.A. Wingens, Ruiqi Liu, Leo A.B. Joosten, Johan Bussink, Klaas W. Mulder

## Abstract

Intratumor heterogeneity negatively impacts therapeutic response and patient prognosis. Besides the established role of genetic heterogeneity, non-genetic mechanisms of persistence to drug treatment are emerging. Here, we characterise cells selected for their persistence to control, epidermal growth factor inhibition (EGFRi), radiation and combined treatment from low passage head and neck squamous cell carcinoma (HNSCC) cultures. Using a panel of 70 (phospho-)specific DNA-conjugated antibodies we measured activities of 8 signalling pathways, self-renewal, differentiation, DNA damage and cell-cycle, in conjunction with the transcriptional output in single cells, using our RNA and Immuno-Detection (RAID) technology. Six recurrent transcriptional programs reflecting processes including proliferation, differentiation and metabolic activity, as well as protein-based signalling-states, were associated with drug persistence, while copy number variation inference indicated involvement of non-genetic tolerance mechanisms. Projecting RNA velocity onto the antibody-derived signalling-states suggested a key role for integrin-mediated focal-adhesion signalling in drug-persistence in our cell system. Using machine-learning we derived a core transcriptional signature connected to adhesion-based drug-persistence, which was predictive of poor prognosis in a TGCA HNSCC cohort (hazard-ratio 1.87, p<10-5). Furthermore, functional analyses confirmed that cells expressing high levels of integrin alpha-6 (ITGA6) were tolerant to EGFRi treatment, and that forcing cells out of this cell-state through transient targeted inhibition of Focal Adhesion Kinase activity re-instated EGFRi sensitivity in drug persistent cells. Taken together, our single-cell multi-omics analysis identified an actionable adhesion-signalling mediated cell-state driving drug tolerance in HNSCC.

## Introduction

Head and neck squamous cell carcinoma (HNSCC) is a prevalent form of cancer that develops in the mucosal epithelium lining the oral cavity and upper aerodigestive tract, with an estimated 800,000 new diagnoses worldwide each year ^1–3^. Risk factors include tobacco and alcohol consumption, exposure to environmental pollutants, as well as infection with human papillomavirus (HPV) and Epstein-Barr virus (EBV)^1^. Treatment options for HNSCC depend on the location and histopathological disease stage of the primary tumour and other factors^4^. In general, single-modality treatments, including surgery or radiotherapy, are applied for early-stage disease. However, for more advanced stage disease, a combination of surgery with chemo-, radio-, immuno-, or targeted therapy is required^1^. Despite the availability of these therapeutic regimens, treatment of HNSCC remains a challenge and resistance to therapy is a common occurrence ^5^. In such cases, patients either do not respond to the given treatment (i.e. intrinsic resistance), or develop recurrent disease after an initial treatment response (i.e. acquired resistance). Drug tolerant persisters (DTPs) are cells that survive the anti-cancer therapy through pre-existing, or therapy induced resistance mechanisms. DTPs frequently display characteristics of cancer stem cells, including stemness and involvement in epithelial-to-mesenchymal transition (EMT) ^6,7^. Interestingly, some DTPs have been shown to display reversibility of their drug-resistance to regain drug sensitivity upon discontinuation of the treatment (i.e. ‘drug holiday’)^6,7^. Therapy resistance and disease recurrence are associated with the level of intratumor heterogeneity ^8,9,10–12^, which has predominantly been studied at the level of genomic heterogeneity in the context of tumour evolution ^13–15^. Hence drug resistance caused by genetic mutations are well established, yet our understanding of non-genetic drug tolerance mechanisms is still emerging ^16,17^. For instance, in BRAF-mutant melanoma, a subpopulation of cells displaying stochastic and reversible high expression of resistance markers (including high expression of the Epidermal Growth Factor Receptor, EGFR) tolerates BRAF inhibition, and eventually acquires permanent adaptations that make them resistant ^18,19–21^. The EGFR is a well-studied oncogene that influences various mechanisms associated with malignancy ^22,23^. It is overexpressed in 90% of HNSCC cases and this is negatively linked with overall survival ^1,22–25^. Cetuximab is a monoclonal antibody blocking the extracellular domain of EGFR and is FDA-approved for treatment of HNSCC as monotherapy, or in combination with other therapeutic modalities ^1,22,23,26^. In addition to monoclonal antibodies, numerous studies have explored the use of small molecules that act as tyrosine kinase inhibitors to block the intracellular domain of EGFR ^27,28^.

Recent developments in single-cell technologies have provided powerful tools to study cellular heterogeneity in the context of HNSCC biology. For example, single-cell transcriptional profiling (i.e. scRNA-seq) of primary and metastatic HNSCC samples identified cells at the leading-edge of the tumour that undergo a partial EMT(pEMT) are predictive of local metastasis and poor-prognosis associated pathological features ^29^. scRNA-seq analyses of HNSCC samples reflecting disease progression in the form of non-tumoral region surrounding normal lesion, precancerous leukoplakia, primary cancer, and metastatic tumour in lymph node revealed interactions between the tumour, stroma and immune cells in this process ^30^. Prognostic signatures of cancer-associated fibroblasts have also been identified by combining single cell and bulk RNA sequencing ^31^. Moreover, characterization of response to cetuximab treatment in HNSCC cell-lines using single-cell RNA and scATAC sequencing indicated transcriptional and chromatin rewiring as early events during cetuximab treatment^32^.

Despite these advances, the mechanisms giving rise to drug tolerant persister cells in HNSCC, and insights into potential strategies to re-sensitize these, remain unclear. Here, we characterised transcription and signalling states associated with drug tolerant persister cells derived from low-passage patient-derived HNSCC cells in culture using RNA and immunodetection (RAID) ^33,34^. Using this single-cell multi-omics technology to simultaneously quantify RNA expression and intracellular (phospho-)protein levels from individual cells, we identified concordant actionable transcriptional programs and signalling signatures associated with drug tolerance. Moreover, we found that integrin-mediated adhesion signalling is functionally involved in drug tolerance and that transient inhibition of its downstream effector Focal Adhesion Kinase (FAK) restores drug sensitivity in the tolerant population. Taken together, our work reveals the power of single-cell (phospho-)proteogenomic analysis of tumour cell heterogeneity to identify potential routes towards re-sensitization of drug-tolerant persisters.

## Results

### Selection and single-cell multi-omic profiling of drug tolerant persisters from cultured HNSCC cells

To identify potential mechanisms leading to drug tolerance, we sought to enrich cells from HNSCC cultures that tolerated 2 weeks of treatment with epidermal growth factor inhibitor (EGFRi), a dose of radiation, or a combination of radiation with EGFRi treatment (Figure 1a). Non-treated cells were cultured in parallel as a control group. To investigate inherent differences in molecular processes in these cells, rather than the direct effect of the treatment, the cells were allowed to recover for 1 week in the absence of external selective pressure. The resulting populations were then subjected to cell-biological characterisation and single-cell molecular profiling with RNA and ImmunoDetection (RAID). This technology allows us to measure signalling pathway activities, cell cycle, self-renewal and differentiation by quantifying the levels of ∼70 intracellular (phospho-)proteins with DNA-barcoded antibodies, as well as the transcriptome from individual cells to gain insight into distinguishing processes between selected drug-tolerant and control cells (Figure 1a).

To identify appropriate lines for our study, we analysed publicly available exome-sequencing data of 16 low-passage (< passage 12) HNSCC cultures ^35^ and separated these lines into 3 clusters, based on their identified non-synonymous single nucleotide variant profiles (SNVs, Figure S1a, b). Furthermore, these lines could be broadly divided into 4 classes based on their variable allele frequency (VAF) patterns (Figure S1c). Lines SJG26 and SJG17 were selected based on the fact that they belong to two distinct VAF classes and showed non-overlapping SNV profiles (Figure S1a). Additionally, SJG17 has a high genomic mutation burden with many mutations and a VAF of between 0.3-0.6, whereas SJG26 harbours fewer mutations with low VAFs, indicating differences in genetic heterogeneity within these cultures (Figure S1d). Based on dosing experiments that included control non-cancer keratinocytes (Figure S1e), these lines were treated with vehicle (NT), or were subjected to treatment with EGFRi (AG, 2.5 µM AG1478), radiation (R, single dose of 8 Gy) or EGFRi combined with radiation (RAG, 1.25 µM AG1478 + 4 Gy) at the time-points indicated in Figure 1a. Following the 1-week recovery period, the selected cells were re-challenged with the original treatment, showing that the selected populations from the SJG26 line largely retained their tolerant phenotype (i.e. semi-stable tolerance), while the SJG17 derived cells reverted back to a drug sensitive state (i.e. transient tolerance, Figure 1b). Tracking cell growth using live-cell imaging indicated that the selected cell populations grew at rates similar to the controls, with the exception of the R and RAG selected population from the SJG17 line, which showed somewhat slower growth (Figure 1c). Moreover, the growth-rate of all selected populations was reduced upon retreatment with 5 µM AG1478, indicating residual drug response following the recovery period (Figure 1d). Thus, we established selection of drug-persistent cell populations from two low-passage HNSCC lines.

**Figure 1:**
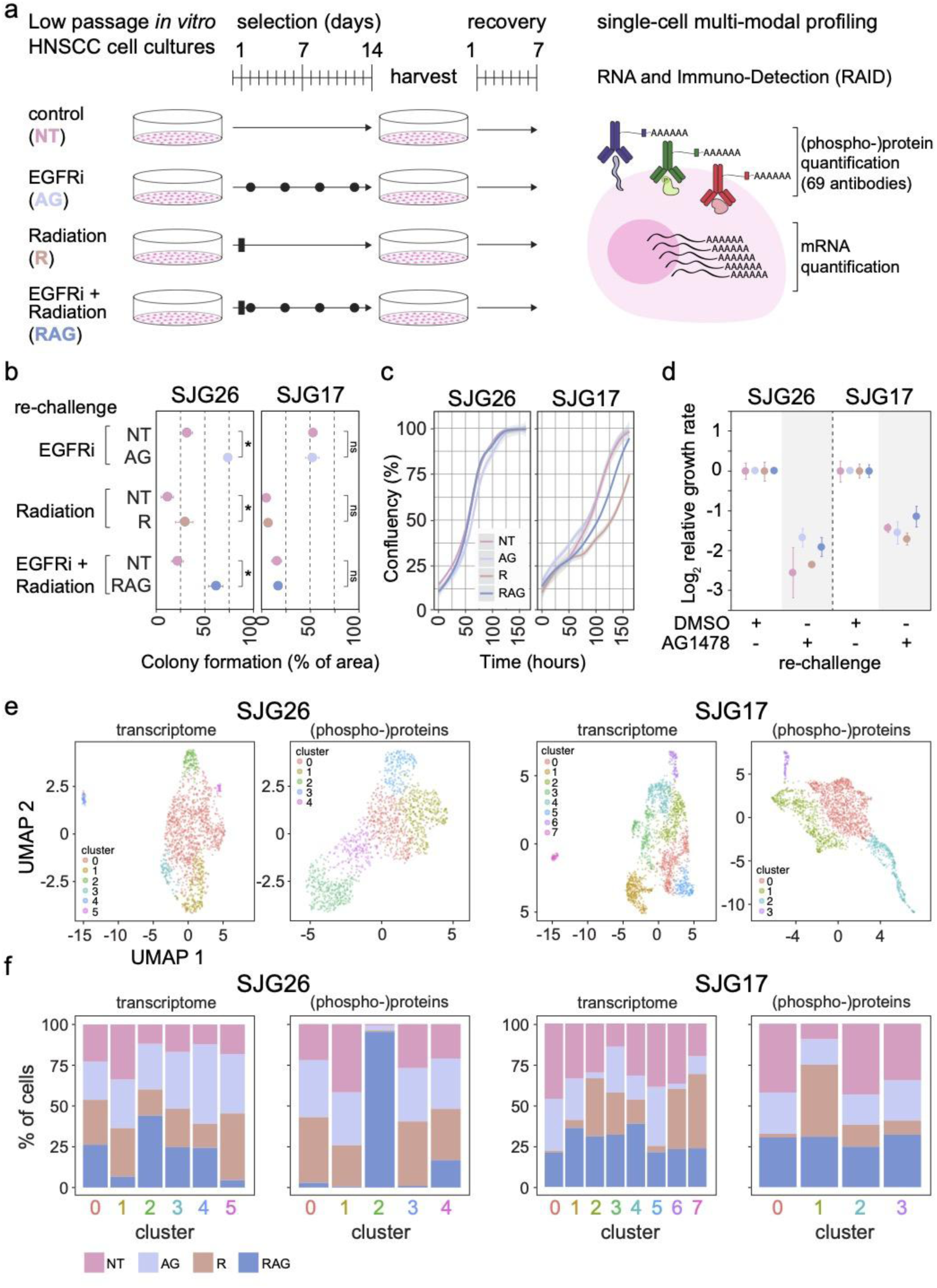
Selection and single-cell multi-omics profiling of drug-tolerant persisters from cultured HNSCC cells. **(a)** Schematic representation of the experimental design to select drug-tolerant cells. Closed circles and rectangles indicate treatment timing. Medium was refreshed at the same time as AG1478 treatment for all cultures. **(b)** Colony formation assay of re-treated drug-tolerant populations from SJG26 and SJG17 HNSCC lines, represented as quantified percentage of covered area of the culture plate. n=3, error bars denote SD, * indicates p<0.05, two-tailed t-test. **(c)** Growth curves (% confluency of the plate) of selected populations from SJG26 and SJG17 HNSCC lines, under standard culturing conditions, derived from live-imaging for 7 days. Curves derived with Loess regression, shaded area indicates 95% CI. **(d)** Relative growth-rates calculated from the linear part of live-imaging based growth-curves of selected drug-tolerant cells from SJG26 and SJG17 HNSCC lines with or without continuous (re-)treatment with AG1478 (5mM). n=3, error bars denote SD. **(e)** UMAP representations of selected drug-tolerant cells from SJG26 or SJG17 HNSCC lines based on either RNA transcriptome or antibody-derived (phospho-)protein measurements. Colours indicate clusters identified using Leiden clustering. **(f)** Proportions of cells from the different selected populations from SJG26 and SJG17 HNSCC lines for each identified transcriptome and (phospho-)protein data derived clusters, respectively.

To characterise potential transcriptomic and (phospho-)proteomic subpopulation structures, we subjected the selected populations to single-cell RAID profiling. We used our previously validated panel of 70 antibodies covering 9 key developmental and cancer-associated signalling pathways (EGFR, Wnt, Notch, TGFβ, TNFα, JAK-STAT, NFκB, BMP and Integrin-mediated adhesion), as well as cell-cycle/proliferation, self-renewal and differentiation processes (Table S1)^36,37^. Over half of the included antibodies were directed against phosphorylated epitopes to measure signalling activity. To allow intracellular antibody staining, the cells were fixed using chemically reversible cross-linkers followed by permeabilization, blocking, antibody staining, washing and distribution as individual cells into 384-well plates containing reverse cross-linking buffer ^33^. We adopted a modified SORT-seq/CEL-seq2 work-flow to simultaneously quantify the transcriptome and antibody derived tags (ADTs) for each individual cell ^33^. This resulted in median detection of 1550-2100 genes per cell (including only cells with >500 genes detected), resulting in a total of 1440 and 2067 cells from SJG26 and SJG17, respectively (Figure S2a, b). We detected > 90% of the antibodies in > 90% of the cells (Figure S2a, b) with a median of 25-85 counts/antibody per cell (range 0-325, depending on the antibody). Following normalisation and batch-correction (see methods section for details), these data were used to call clusters and represented in UMAPs based on variable genes (Figure 1e, transcriptome) or antibodies (Figure 1e, (phospho-)proteins)). This revealed multiple population sub-structures based on the transcriptomic and proteomic data for the SJG26 and SJG17 lines, respectively. Cells derived from the different drug-tolerant selected populations (NT, AG, R and RAG) were relatively well distributed over the different transcriptomic clusters (Figure 1f and S2c). In contrast, we observed a strong enrichment of the SJG26 RAG (radiation + EGFRi) and SJG17 R (radiation) persister populations in proteomic clusters 2 and 1, respectively (Figure 1f). These results indicate that we obtained a high-quality multi-omic transcriptomics and (phospho-)protein dataset at single cell resolution to explore potential mechanisms of drug-tolerance in HNSCC.

### Drug persistent populations are not strictly associated with global genetic signatures

As resistance to therapy has been linked to (pre-existing) genetic subpopulations that are favoured under drug-pressure^14^, we investigated whether the persister populations we selected displayed evidence of specific enriched genotypes. First, we inferred single-cell copy number variation (CNVs) from our scRNA-seq data ^33,38^ using data of equal quality and depth of karyotypically normal primary human keratinocytes as a reference ^33^. This approach identified 7 and 8 CNV clusters for the SJG26 and SJG17 populations, respectively (Figure S3a-d). Several clusters contained known clinically relevant karyotypic abnormalities, including amplification of chromosome 3q24-26 containing the *PIK3CA* gene (SJG26 CNV cluster 3) and deletion of sections of chromosome 11 harbouring the *FADD* and *YAP1* genes ^39^. Notably, all clusters contained cells from the control and all selected populations, indicating that there was no absolute co-segregation of inferred CNV genotype and persister phenotype (Figure S3e, f). Second, we examined variably expressed SNVs from bulk RNA-seq as a measure of enrichment for genetic subpopulations. For this, we specifically searched for the known SNVs identified from the available exome sequencing data in the SJG17 (118 detected with > 20 reads/sample) and SJG26 (13 detected) lines ^35^ (Figure S3g, Table S2). Several genes in the SJG17 line contained multiple SNVs in the population and showed concordant VAFs in the different persister populations, suggesting they occurred in the same subpopulation/cells and validating the approach (Figure S3h). We observed some enrichment for SNVs in the radiation selected SJG17 persister population, but not the other SJG17 and SJG26 populations (cut-off p < 0.01, log_2_FC > 2, Figure S3i, j). Notably, the SJG17 line showed a transient tolerance (Figure 1b), whereas a true genetically driven drug tolerance is expected to be stable upon selection. Although these analyses suggest a contribution of at least some potential genetic component to drug tolerance in the SJG26 and SJG17, there were no dominant global transcriptional and genetic profiles strongly associated with drug persistence in the selected populations. In contrast, some of these populations could be clearly distinguished based on their (phospho-)protein expression phenotypes (Figure 1e), suggesting that differences in signalling activities may play an important role in persisting drug treatment.

### Identification of drug-tolerance associated RNA expression programs

To identify potential biological processes active in subpopulations of persister and control cells, we defined differentially expressed genes from their single-cell transcriptional profiles using two independent approaches. First, we used non-parametric statistical tests (Wilcoxon Rank Sum) to find marker genes for the clusters present in the two datasets using the Seurat framework (Table S3)^40^. Second, we used a cluster-agnostic non-negative matrix factorization (NMF) approach to derive programs of co-expressed genes in subsets of cells as previously described (Table S4)^29,41^. The resulting cluster-based marker genes and NMF programs were then assessed for overlapping genes, revealing high concordance between the two approaches (Figure 2a). Next, these programs were collapsed into 6 robust expression programs that were interpreted and annotated using Gene Ontology over-representation analysis (Figure 2a, b, Table S5). Assigning a program score (derived using a summation approach) to each individual cell confirmed that these 6 distinct transcriptional programs were active in cells derived from both independent HNSCC lines (Figure 2a, c). Moreover, these programs were differentially active in the various selected persister populations and between the different lines. For instance, several of the persister populations were associated with higher proliferation/cell-cycle program scores and a concomitant increase in cellular respiration score (Figure 2d). Conversely, the same populations displayed decreased expression of the epithelial differentiation program (Figure 2d), suggesting a potential role for cellular proliferation, differentiation, as well as cellular respiration/metabolism in drug tolerance. As EGFR activity is a main factor in HNSCC, we investigated whether the identified robust expression programs were dependent on EGFR pathway activity. We subjected control (NT) and EGFRi (AG) selected persister cells from both SJG17 and SJG26 to short-term treatment with the EGFR inhibitor (5 µM AG1478 for 3 hours) followed by RAID profiling. This revealed that sustained EGFR signalling modulates these robust expression programs (Figure S4a and b). Moreover, cell line and drug-persister population dependent effects suggest a rearrangement in the regulation of these programs after, or during, selection of drug-tolerant cells. We interrogated if these different cell-states might be connected and whether some may transition into another using RNA velocity analysis. For this, the same set of variable genes used to identify the six transcriptional programs were used to derive velocity estimates with scVelo^42^. For both the SJG17 and SJG26 lines we noted that RNA velocity (visualised as streamlines on the RNA UMAP coordinates) tends to flow from the cells with a high score for the cellular respiration RNA program to cells with high proliferation/cell-cycle program scores and finally to the cells expressing high levels of epithelial differentiation genes (Figure S4c and Figure 2c). This suggests that cells may transition between cell- and transcriptional states, and that cells in various states may display differences in drug sensitivity and tolerance. In summary, the transcriptome component of our RAID profiling identified 6 robust RNA expression programs and cellular processes associated with drug-persistence in both HNSCC lines.

**Figure 2:**
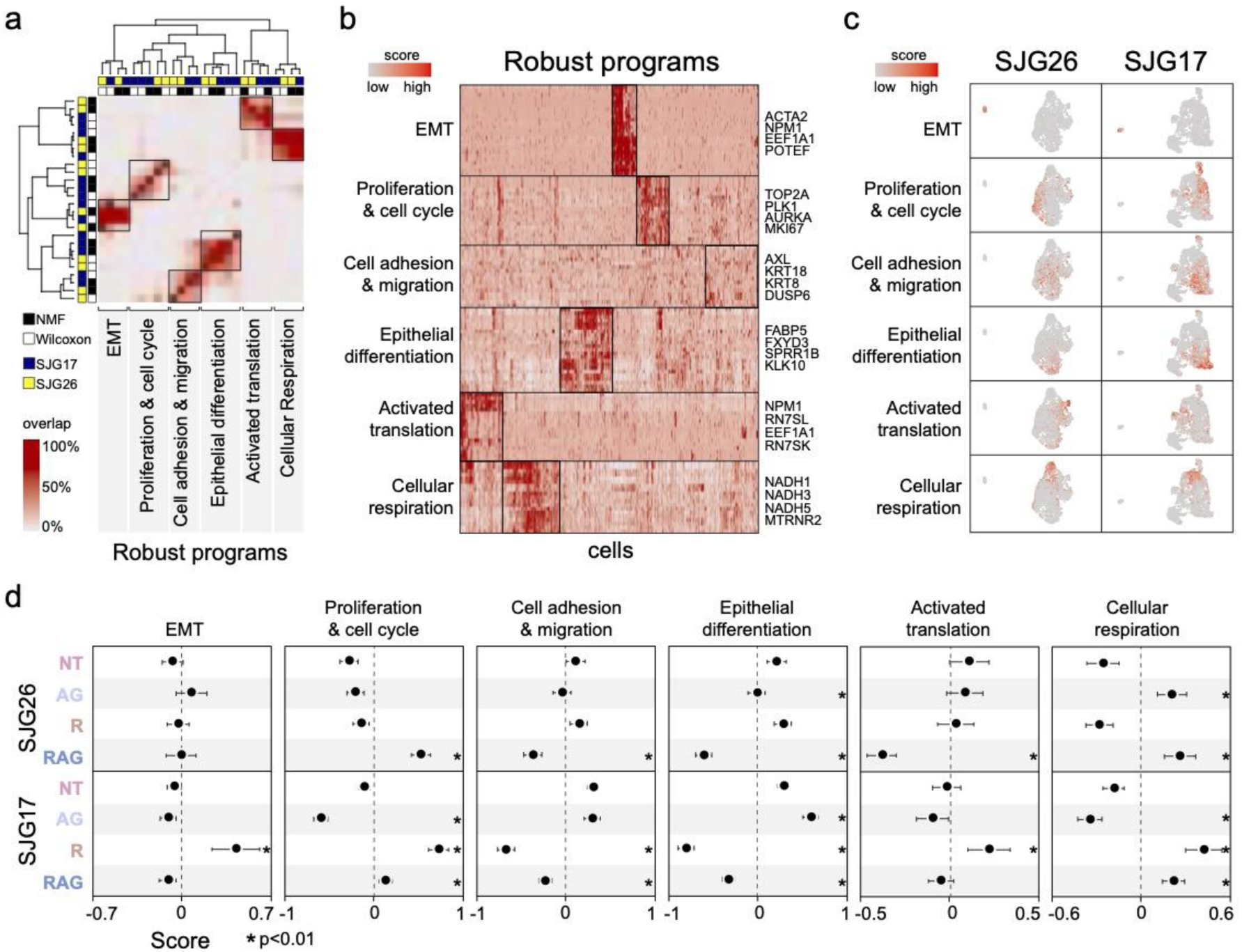
Identification of drug-tolerance associated RNA expression programs. **(a)** Heatmap displaying the overlap between the identified differentially expressed genes per cluster (Wilcoxon rank sum test q<0.05) and Non-negative Matrix Factorization (NMF) derived cluster independent gene programs in the SJG26 and SJG17 datasets. Programs clustering together (hierarchical clustering) were joined into so-called Robust programs, and subjected to GO-enrichment analysis for biological interpretation. **(b)**Clustered Heatmap of normalised expression of the top25 genes per Robust program in a maximum of 100 cells per cluster from SJG26 and SJG17, respectively. Example genes are indicated on the right for each program. **(c)** Transcriptome based UMAP representation of SJG26 and SJG17 datasets with superimposed normalised robust program scores. **(d)** Robust expression programs are enriched various selected drug-tolerant populations from both SJG26 and SJG17 HNSCC lines. Forest plots display the average program score for each indicated population. Error bars denote 95% CI, * indicates p<0.01, two-sided t-test compared to NT.

### Identification of drug-tolerance associated intracellular signalling activities

A unique and powerful feature of RAID is the quantification of the transcriptome in conjunction with many intracellular phosphorylated proteins for each cell. Our validated panel of DNA-conjugated antibodies measures ∼70 components of key cancer-related signalling pathways (EGFR, Wnt, Notch, TGFβ, TNFα, JAK-STAT, NFkB, BMP and Integrin-mediated adhesion), as well as cell-cycle regulators and epithelial stem- and differentiation markers ^36,37,43^. Statistical analyses revealed differentially expressed (phospho-)proteins in the selected drug-persister populations (AG, R and RAG) versus control (NT) for both SJG17 and SJG26 lines (Figure 3a, Table S6). We visualised these (phospho-)proteins in the context of their cognate biochemical pathways (Figure S5a), with node colour and size representing the log_2_ fold-change and −log_10_ p-value (Kolmogorov-Smirnov test) of the comparison to the control (NT) population, respectively (Figure 3b). This highlighted the most significantly affected nodes and revealed that the measured proteins within a pathway generally behaved concordantly (Figure 3b and S5b, c). To investigate activity at the pathway level, rather than isolated nodes, we derived a summarising score for each pathway for each individual cell (see Methods section and Table S7 for details). This revealed significantly increased activity of Integrin-mediated adhesion and decreased Notch signalling in the SJG26 RAG and SJG17 R and RAG persister populations (Figure 3c). Superimposition of these scores for each cell onto the protein UMAPs showed differential and gradual transitions in pathway activities between regions of the UMAP, as well as in the clusters defined using the (phospho-)protein measurements in both the SJG26 (Figure 3d and Figure S6a, b) and SJG17 (Figure S6c, d) lines. We noted that SJG26 cluster 2 displayed the highest scores for adhesion signalling, with cluster 4 showing an intermediate level of activity (Figure 3d). In contrast, other pathways including JAK-STAT and Notch were comparatively low in cells from these clusters (Figure 3d). Examples of antibodies representing these pathways are displayed in Figure 3d. Taken together, these results show that the selected persister populations differ in their pathway activities, suggesting the existence of a potential drug-tolerant signalling phenotype.

**Figure 3:**
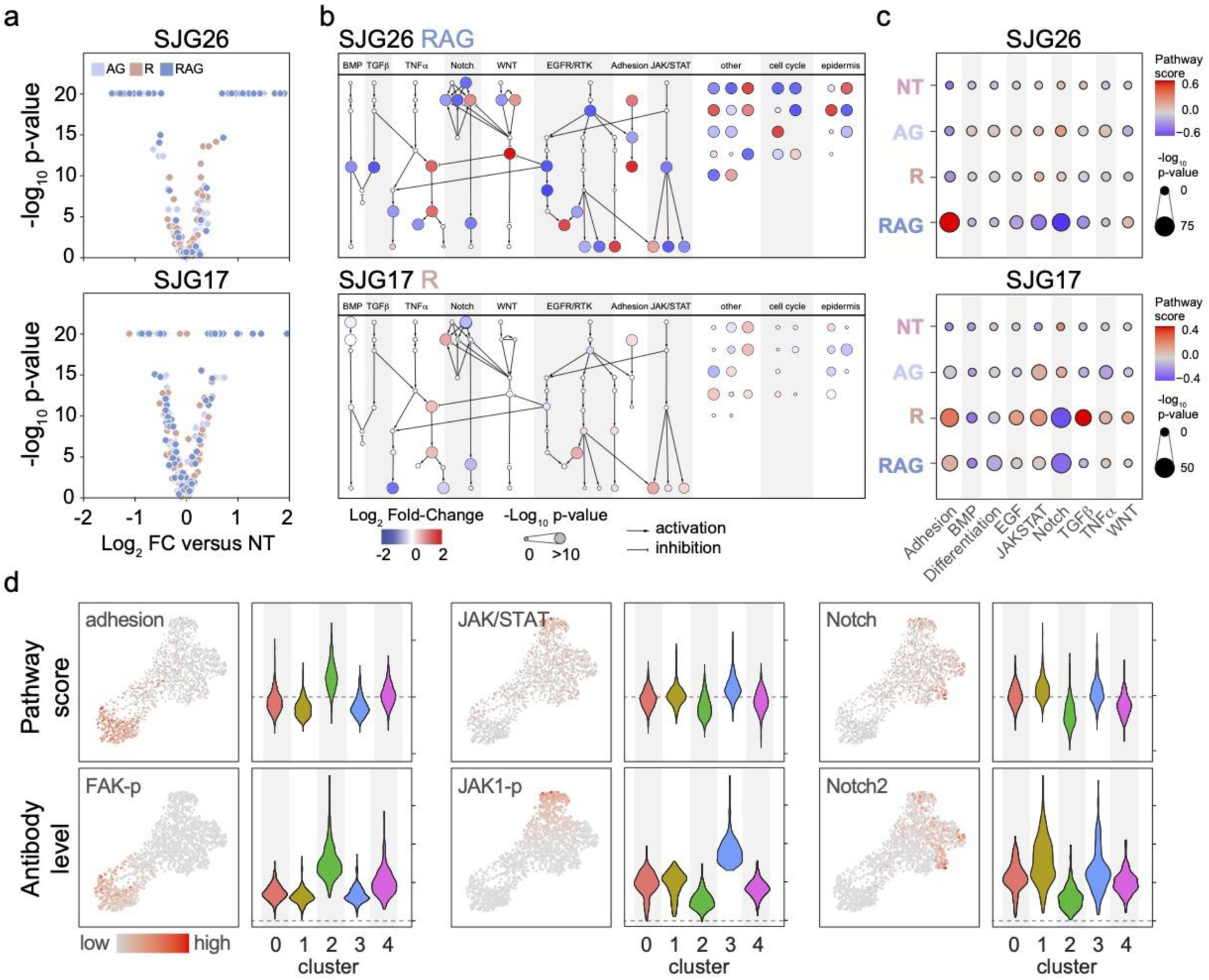
Identification of drug-tolerance associated intracellular signalling activities. **(a)** Volcano-plots of differential single-cell antibody-signal testing (Kolmogorov-Smirnov (KS) test) in the drug-tolerant populations to the NT control cells in SJG26 and SJG17 HNSCC lines, respectively. **(b)** Differential abundance analysis superimposed on a biochemical network of signaling pathways and processes covered by the RAID antibody panel. Colour indicates the effect size and the node size represents its statistical significance (-log_10_ p-value, KS-test). Arrows depict known kinase-substrate pairs. Identity of the nodes is depicted in Supplemental Figure 5A. **(c)** Dotplots of normalized (z-score) pathway scores drug-tolerant populations from SJG26 and SJG17 HNSCC lines, respectively. Colour indicates the effect size and the dot size represents its statistical significance (-log_10_ p-value, two-sided t-test) compared to the NT control population. **(d)** Selected pathway scores (Adhesion, JAK/STAT and Notch) per individual cell superimposed on the (phospho-)protein UMAPs for the SJG25 HNSCC line (top row) and a representative antibody signal for these pathways (bottom row). Violin plots represent the signal distribution of the pathway scores (top) and antibody levels (bottom) per (phospho-)protein-based cluster, respectively. Violin plot colours indicate clusters as called in Figure 1e.

### Identification of a drug-persister signalling phenotype associated with cellular respiration

We wondered whether the complex signalling gradients we observed were likewise reflected in the transcriptome of the cells and whether this may further inform us about cell- and signalling-state transitions involved in drug-tolerance. We made use of the fact that we have fully matched transcriptome and protein data for each cell and projected the RNA velocity (as calculated for Figure S4c) onto the (phospho-)protein UMAP. This revealed that the direction of the RNA velocity in the SGJ26 line diverges at the boundary between clusters 0 and 4 in the SJG26 line (Figures 4a and S2d). Conversely, the RNA velocity streamlines in the SJG17 line point towards the region of the UMAP occupied by the control (NT) population (Figures 4a and S2c). These observations suggest that the SJG26 derived cells represent two (semi-)stable states, whereas the SJG17 cells will revert back to a single state. This notion is consistent with our observation that the drug-tolerant persister populations selected from the parental SJG26 largely retain their tolerant phenotype after the 1-week recovery, whereas those selected from SJG17 revert their drug-tolerance and retain their drug-sensitivity (Figure 1b).

**Figure 4:**
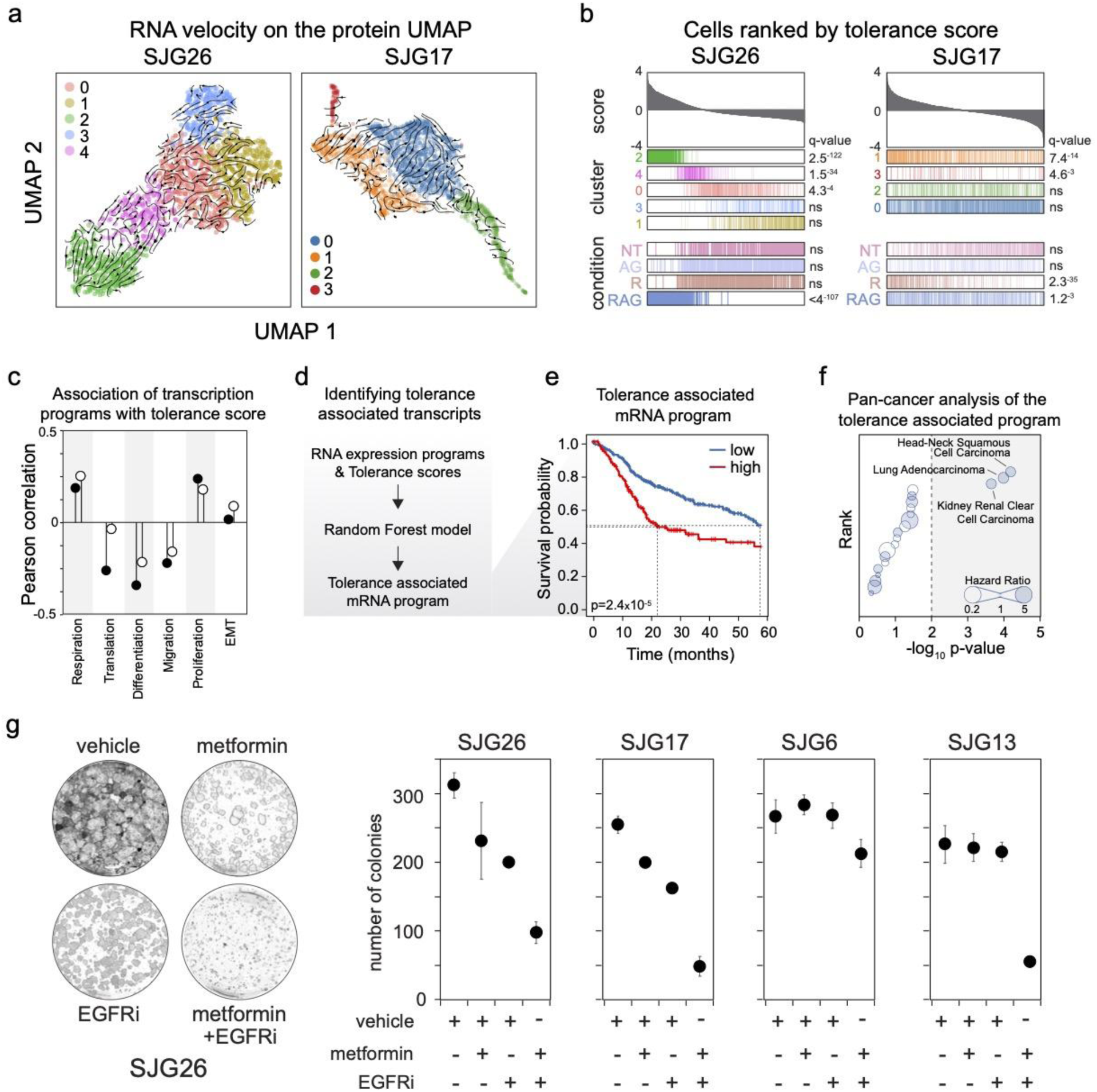
Identification of drug-tolerance signalling phenotype associated with cellular respiration. **(a)** RNA velocity projected onto the (phospho-)protein derived UMAPs. Colours indicate clusters as called in Figure 1e. Arrows indicate direction of RNA velocity. **(b)** Rug plots representing the distribution of cells from the indicated clusters and drug-tolerant populations after ranking on the calculated tolerance score for the SJG26 and SJG17 HNSCC lines, respectively. q-values indicate significance (KS-test) of their enrichment among high tolerance-score cells. ns denotes non-significant. **(c)** Pearson correlation between the tolerance score and the robust program scores across all cells in the SJG26 (filled circle) and SJG17 (open circle) HNSCC lines, respectively. **(d)** Work-flow to derive a Random Forest machine-learning based tolerance score-associated mRNA signature. All cells from both SJG26 and SJG17 HNSCC lines were used to derive the mRNA signature. **(e)** Kaplan-Meier analysis of Overall Survival of patients in the TGCA HNSCC cohort separated by high (red line) or low (black line) expression of the tolerance-associated mRNA signature. Log-rank p-value at 1% FDR. Dotted lines indicate median survival (0.5 survival probability). **(f)**Summarised overall survival analysis of patients with high vs low expression of the tolerance-associated mRNA signature across TCGA cancer-type cohorts. Results are ranked on −log_10_ p-value with the data point size representing the Hazard-Ratio (HR). The dotted line indicates a p<0.01 significance cut-off. **(g)** Colony formation assays (CFA) of control, metformin, EGFRi or metformin and EGFRi co-treatment of the indicated parental SJG HNSCC lines. Left: representative images of 1 replicate of CFA for SJG26. Right: quantified results from the indicated lines. N=3, error bars denote SD. Controls were vehicle (DMSO) treated at the same concentration as the drug-treatments.

The RNA velocity results combined with the observed gradients of signalling activities across the UMAP of the SJG26 cells, suggest that the cells in cluster 2 may exhibit a drug-persistent signalling phenotype. As such, the combined activities of the (phospho-)protein features distinguishing these cells might represent a drug tolerance state. To explore this hypothesis, we constructed a ‘tolerance score’ for each cell based on the cluster 2 specific protein markers (see Methods section and Table S7 for details). Ranking the cells by this score showed that SJG26 cells from clusters 2, 4, 0 and RAG selected cells display significantly higher scores, as expected (Kolmogorov-Smirnov test, q<0.01, Figure 4b left panel and Figure S7a), consistent with the gradients of signalling activities observed in these clusters (Figures 3d and S6a). Importantly, calculating the tolerance score for the SJG17 cells revealed that the SJG17 R and RAG selected populations also exhibited significantly higher scores (Figure 4b right panel and Figure S7b). This indicates that the potential signalling phenotype captured by the tolerance score, even though it was solely based on the SJG26 data, may be indicative of tolerance in both independently derived sets of lines. In agreement with this notion, the global RNA velocity directions closely align with this tolerance score in both datasets (Figure S7c), suggesting a coupling of the signalling phenotype with gene expression. Indeed, the tolerance score displayed a consistent positive correlation with the respiration and proliferation RNA expression programs, and a negative correlation with the differentiation and migration expression programs in both SJG26 and SJG17 (Figure 4c and Figure S8a,b). This suggests that the signalling phenotype and gene expression programs we identified are jointly connected to drug tolerance.

To gain insight into which of the genes represented in the 6 robust expression programs were most tightly associated with the tolerance score, we applied Random Forest (RF) Regression on the union of their most robustly expressed genes in SJG17 and SJG26 (216 genes in total, see Methods section and Table S8 for details), using the tolerance score as the response variable. This resulted in a model that explained 52% of the variance with a mean of squared residuals of 0.11 from 1000 random tree initiations. The importance for each gene in this model was quantified by the percentage increase in mean squared error (%IncMSE) of the prediction when the gene was left out (also called mean decrease in accuracy, Figure S8c). We calculated an empirical p-value of this importance for each gene through randomisation, identifying sixty candidates that displayed a higher than the average importance and p < 0.01 (Figures 4d, S8c and Table X). As an orthogonal confirmation approach, we applied step-wise linear regression based on the Akaike Information Criterion (AIC) using the same set of transcribed genes as input and the tolerance score as the dependent variable. The resulting model (R^2^ = 0.59, p = 2.2*10^-16^) comprised 124 of the 216 input genes, including > 70% of the 60 important and significant genes that were identified by the RF model, providing confidence in our approach to identify tolerance-associated RNA transcripts.

If the expression of these genes is indeed indicative of tolerance to treatment, we would predict them to also show an association with HNSCC patient survival in real world data. We explored this using the publicly available data from The Cancer Genome Atlas (TCGA). In total, 41 of the 60 tolerance associated transcripts were reliably detected in the TCGA Head and Neck cancer RNA-seq cohort^44^ and 18 of these showed significant association with poor patient prognosis (Figure S8d and Table S9). When combined, these genes represent a tolerance associated RNA program that is predictive of patient overall survival (Figure 4e, p < 1*10^-5^, 1% FDR and hazard-ratio = 1.87). Moreover, this program is associated with overall survival in an additional 2 out of 21 cancer-types annotated in the TCGA database, indicating that the tolerance associated gene expression we identified reflects cancer-type specific, rather than general mechanisms (Figure 4f). Notably, several mitochondrially encoded transcripts that were part of the respiration expression program were predictive of poor prognosis on their own and were differentially expressed in cell clusters displaying an increased tolerance-score (Figures S8d, e).

To explore this association between drug-tolerance and energy metabolism further, we measured real-time oxygen consumption rates (OCR, to measure respiration) and extracellular acidification rates (ECAR, to measure glycolysis) of the selected persister cell populations (Figure S9a,b). Indeed, basal respiration was elevated in the SJG26 AG and R, as well as in the SJG17 R and RAG populations (Figure S9c, d), consistent with a role for respiration in drug-tolerance. When treated with the mitochondrial respiratory chain complex I inhibitor metformin, the SJG17 and SJG26 persister populations showed differential dependence on mitochondrial respiration for their growth-rate (Figure S9e). Additionally, maximum respiration capacity and glycolysis were affected in multiple persister populations, further supporting the notion that altered mitochondrial energy metabolism is linked to drug-tolerance in these HNSCC lines (Figure S9a,b). To functionally test this, we performed clonal growth assays in the presence of vehicle, EGFRi, metformin, and the combination of both drugs. Combined treatment with these inhibitors indeed prevented the emergence of EGFRi persistent cell populations in both SJG17 and SJG26 lines, functionally implicating mitochondrial energy metabolism in drug tolerance in our system (Figure 4g), in line with previous findings that taking metformin was associated with improved overall survival in HNSCC patients ^45^. Taken together, we identified a drug-tolerance signalling-phenotype and its associated transcriptional program involving mitochondrial energy metabolism. Moreover, we functionally implicate this cellular respiration in tolerance to EGFR inhibition in vitro and tolerance-associated gene expression with poor patient prognosis in a clinical cohort.

### Transient inhibition of Focal Adhesion signalling reinstates drug-sensitivity

With establishing that this tolerance score is indeed informative and representative of a functional and actionable drug-tolerant cell-state, we should be able to identify markers and mechanistic mediators of drug-tolerance by identifying the contributing biochemical pathways. To this end, we calculated the correlation between the tolerance score and individual signalling pathway activity scores covered by our antibody-panel, highlighting three pathways that showed consistent associations for both HNSCC lines. We found a strong positive correlation for adhesion signalling (R = 0.82 and R = 0.67 for SJG26 and SJG17 respectively) and a weaker positive correlation with Wnt pathway activity (R = 0.21 and R = 0.26 for SJG26 and SJG17 respectively, Figure 5a, b and Figure S10a). In contrast, we found a strong to moderate negative correlation with Notch pathway activity (R = −0.59 and R = −0.28 for SJG26 and SJG17 respectively, Figure 5a, b). Activated adhesion signalling is found in the basal layer of the normal human epidermis containing the proliferative progenitor/stem cells that are anchored to the underlying basement membrane, whereas Notch signalling is active in cells as they form the suprabasal differentiated layers in normal human epidermis ^46^. This suggests that drug persister cells harbour some signalling features of progenitor/stem cells that may be involved in establishing or maintaining their drug-tolerant phenotype. To solidify these findings and test the functional involvement of adhesion signalling in drug-tolerance, we sought to prospectively isolate potential persister cells from the parental cell lines. We noted that the cell surface adhesion molecules Integrin alpha-6 (ITGA6, also called CD49f) and Integrin beta-1 (ITGB1) were among the proteins contributing to the tolerance score (Figure 5c, d). We stained the parental SJG26 cells with an ITGA6 antibody and isolated live ITGA6^low^ and ITGA6^high^ cells using fluorescent activated cell sorting. These cells were seeded onto mitotically inactivated mouse feeder cells and allowed to attach and recover. Two days after seeding the cells were treated with 0, 2.5 or 5 µM EGFR inhibitor (AG1478) continuously for 10 days, after which the plates were fixed and the colonies visualised and quantified. We found a dose-dependent decrease of colony size, with the ITGA6^high^ cells giving rise to significantly larger colonies (Figures 5e and S10b). This result indicates that ITGA6^high^ cells, representing cells with an increased tolerance score, indeed reside in a pre-existing drug-persistent state. Moreover, this state may be maintained by ongoing intracellular signalling initiated by integrin-mediated adhesion, as reflected by increased Focal Adhesion Kinase (FAK) phosphorylation in the top 10% cells with the highest tolerance scores (Figures 5f and S10c).

**Figure 5:**
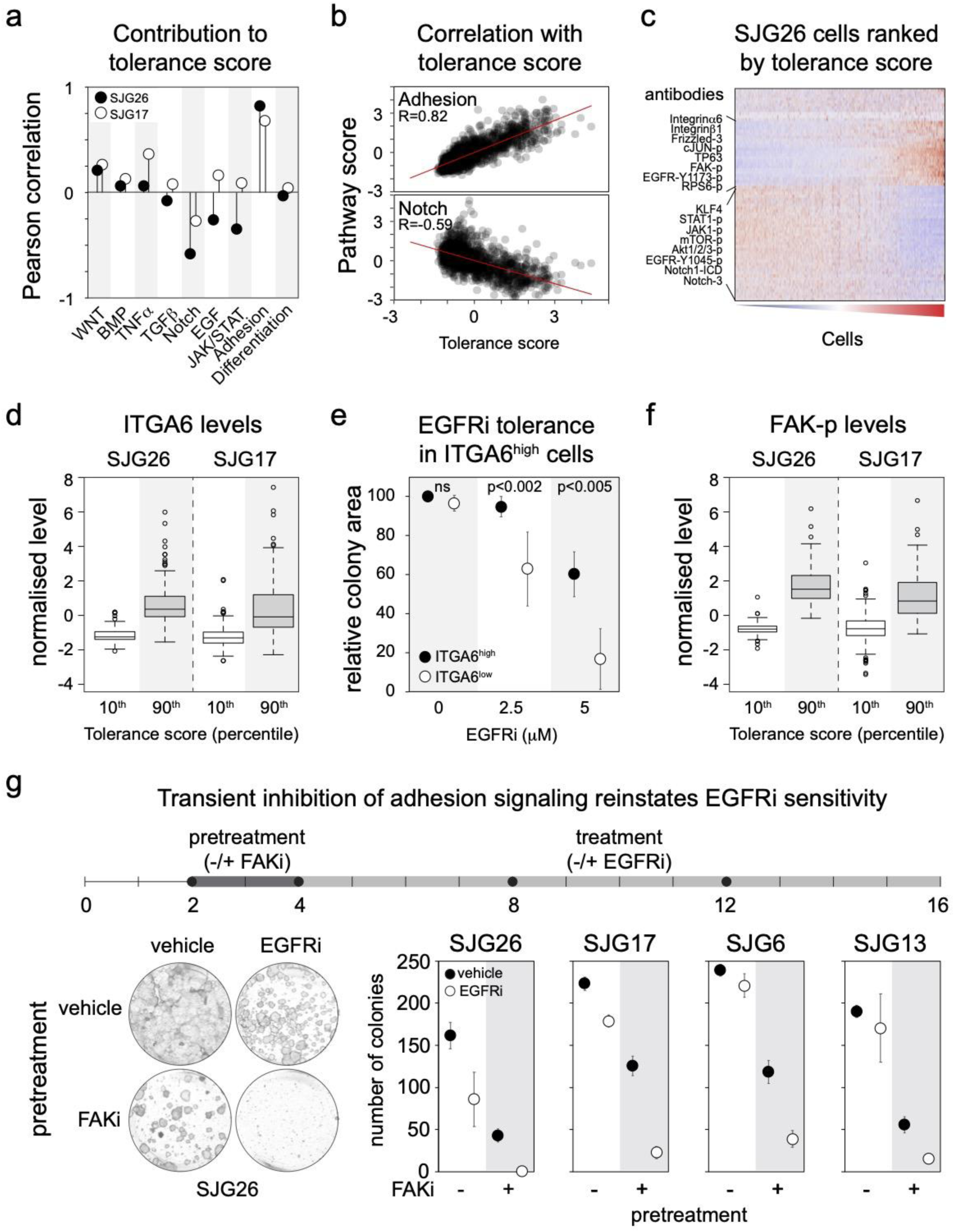
Transient inhibition of Focal Adhesion signalling reinstates drug-sensitivity. **(a)** Pearson correlation between the tolerance score and the indicated pathway scores across all cells in the SJG26 (filled circle) and SJG17 (open circle) HNSCC lines, respectively. **(b)** Scatter plots of the tolerance score versus adhesion and Notch pathway scores in SJG26 cells, respectively. R indicates Pearson correlation coefficient. Red line represents linear regression on the data. **(c)** Heatmap of normalised levels of all measured (phospho-)proteins from the SGJ26 dataset. Data is clustered on the antibody signals and cells are ranked by their tolerance score (low to high). Example (phospho-)proteins positively or negatively correlated with the tolerance score are indicated on the left of the heatmap. **(d)** Box-plot representing the ITGA6 levels of the 10^th^ and 90^th^ percentile of cells based on their tolerance score in the SJG26 and SJG17 datasets, respectively. **(e)** Colony formation assay of SJG26 cells prospectively isolated using FACS based on low versus high ITGA6 cell surface expression levels, respectively. Cells were allowed to attach and recover for 48 hours before starting treatment with the indicated EGFRi concentrations for 12 days and quantified as percentage of covered area of the culture plate. Point-plots indicate average of n=3, error bars denote SD. P-values for comparison of ITGA6 low versus high results for each EGFRi concentration are indicated (two-sided t-test). **(f)** Box-plot representing the phosphorylated Focal Adhesion Kinase (FAK-p) levels of the 10^th^ and 90^th^ percentile of cells based on their tolerance score in the SJG26 and SJG17 datasets, respectively. **(g)** Colony formation assay of SJG26, SJG17, SJG6 and SGJ13 HNSCC lines pre-treated with vehicle or FAK inhibitor for 48 hours, followed by vehicle or EGFRi treatment (5 mM) for 14 days. Top panel represents the experimental time-line. Closed circles indicate medium changes, numbers indicate days. Bottom left panel, representative images of colonies that formed (SJG26 line) in the indicated conditions. Bottom right panel, point-plots indicate average number of colonies formed in the indicated conditions. N=3, error bars denote SD.

We aimed to coax the pre-existing drug-tolerant persister cells in our parental HNSCC cultures out of this state to re-enter a drug-sensitive state by modulating adhesion signalling. To achieve this, we transiently inhibited FAK (a key downstream effector of integrin signalling) with a well-described small-molecule inhibitor (48 hours, 5µM PF-573228/FAKi) prior to treating the cells with EGFR inhibitor (2.5 µM, 12 days in the absence of FAKi), followed by visualisation and quantification of the colonies that formed (Figure 5g). Conditions without FAKi pretreatment or EGFRi were included as controls. In line with our prediction, drug-tolerant persister populations failed to emerge from the parental SJG26 and SJG17 HNSCC lines after the FAKi pretreatment (Figure 5g). These results suggest that ongoing Focal Adhesion Kinase signalling is required to maintain the drug-tolerant population and that moving cells out of this state reinstates drug sensitivity.

Taken together, our results establish a functional role for Integrin-mediated adhesion signalling in drug persistence, and that targeted and transient inhibition of FAK activity reinstates sensitivity to EGF receptor blocking drugs.

## Discussion

In this study we applied our previously developed single-cell RNA and ImmunoDetection (RAID) technology to simultaneously measure the transcriptome and ∼70 intracellular (phospho-)proteins to identify potential drug persistence mechanisms in cultured head and neck squamous cell carcinoma (HNSCC) cells. Our experimental design allowed us to study resistance over relatively short treatment and recovery periods to capture the role of dynamic processes, including cellular signalling. These experiments were performed on two low-passage HNSCC lines derived from primary patient material (passage < 12) with distinct mutational profiles and burdens to ensure heterogeneity in the parental population. Despite this genetic variation, we failed to find a dominant association of drug persistence and pre-existing genetic subclones, indicating the involvement of orthogonal underlying processes. Our single-cell multi-omics data analyses highlighted multiple potential cellular processes, including proliferation, differentiation, energy metabolism and several signalling pathways associated with drug sensitivity. Combining the identified transcriptional programs and signalling activities in the various isolated drug-tolerant populations, with machine learning revealed an RNA signature associated with tolerance *in vitro* and patient outcome in a clinical cohort. Analysis of this signature indicated involvement of aberrant mitochondrial respiration in drug sensitivity, which was confirmed experimentally. Mechanistically, this RNA program was associated with Integrin-mediated adhesion as a potentially actionable pathway, supported by the finding that we prospectively isolate drug-tolerant cells based on differences in their ITGA6/CD49f surface levels. This is in line with previous observations showing ITGA6 as a stemness driver in HPV+ HNSCC expressing the embryonic markers Oct4 and Sox2, as well as displaying tolerance to cisplatin treatment ^47^. ITGA6 is a known stem cell marker in epithelia that contributes to self-renewal, epithelial-to-mesenchymal transition, metastasis and resistance ^48^, whose high expression has been associated with poor prognosis in HNSCC ^49,50^. These results are corroborated by previous findings on the potential of ITGB1 blocking antibodies to sensitise HNSCC cells to radiotherapy ^47,51–53^.

Furthermore, we found that pre-treating the parental cell populations with a FAK inhibitor prevented the emergence of resistant cells, confirming the functional importance for adhesion signalling in drug persistence in our cell system. Although FAKi performed poorly as monotherapy for cancer, it has been observed as a potential therapeutic strategy to overcome adaptive resistance in various cancers ^54^. Phase 1/2 combination trials involving FAKi are currently ongoing for different cancer types, including non-small cell lung carcinoma, pancreatic cancer, mesothelioma ^54^. In regards to HNSCC, FAK has also been observed in other studies as a driver of resistance to radiotherapy and chemotherapy ^55,56^. These observations, together with our data suggest FAK as a potential target to overcome adaptive resistance to radiotherapy, chemotherapy and targeted therapy.

Taken together, our work provides insights into signalling-dependent drug-tolerance mechanisms and connects them to non-genetic tumour heterogeneity as an important feature of therapeutic response. Moreover, our work serves as an example of the power of single-cell multi-omics analyses to provide strategies for rational redirection of drug-tolerant cells into a drug-sensitive state.

## Materials and methods

### Cell culture and selection of drug-persister cells

Squamous cell carcinoma cells used in this study (SJG26, SJG17, SJG6 and SJG13) were a kind gift from Prof Fiona Watt^35^. The cells were cultured on a feeder layer of mitotically inactivated J23T3 cells in complete FAD medium containing 10% FCS as described in^43,57^. This study was conducted in adherence to the tenets of the Declaration of Helsinki. The cells were subjected to three different selection procedures, namely EGFR inhibition, radiation and the combination of both EGFR inhibition and radiation, for a duration of 2 weeks. EGFR inhibition was performed by treating the cells with 2.5 µM AG1478 starting 48 h after seeding. Medium with AG1478 was replaced every 4 days. Radiation with 8 Gy, at a dose rate of 3.1 Gy/min, was performed 24 h after seeding, followed by medium renewal 24 h later and every subsequent 4 days (X-RAD Biological Irradiator; Precision X-ray, Madison USA). For the combination strategy, the cells were treated with 1.25 µM AG1478 and 4 Gy radiation, following the same timing as for the individual selection procedures. We chose lower doses as the combination of 8 Gy radiation with 2.5 µM AG1478 was lethal to all cells during the selection phase.

### Colony formation assays

For the colony formation assays, 1000 cells per well were seeded on a feeder layer of mitotically inactivated J2-3T3 cells in 6-well plates. The treatment regimens matched those used for selection and the colonies were grown for 2 weeks before fixation with 4% paraformaldehyde (15 minutes at RT). The fixed cells were permeabilized with 0.1% triton X-100 in PBS (10 min, RT) and subsequently blocked with 10% bovine serum in PBS for 1 hour at RT. The cells were then stained with anti-ITGβ1 (0.25 ng/µl, DSHB P4C10, 1 hour at RT), washed 3 times with phosphate-buffered saline (PBS, pH 7.4), followed by secondary antibody anti-mouse IRDye 800 (1:4000, LiCor Biosciences) and DR (1:4000, Biostatus) for 1 hour at RT. After washing, the stained plates were imaged using the Odyssey CLx (LiCor Biosciences). The acquired images were quantified by Cell Profiler version 4.1.3. The pipeline used can be found at Github (https://github.com/dwkarjosukarso/HNSCC_manuscript).

### Cell proliferation assay

Cell proliferation assay was performed by seeding 2000 cells on collagen-coated 96-well plate in triplicate, followed by live-cell imaging every 6 h using Incucyte Systems (Sartorius) for the duration of 1 week. Medium was refreshed after 3 days. The confluency of the cells was quantified from acquired images using the Incucyte Analysis software.

### Production of Antibody-ssDNA conjugates

Antibodies were buffer exchanged to 50 mM borate buffer saline (BBS) and subsequently functionalized with NHS-S-S-Tetrazine as previously described, with some modifications^43^. In brief, each antibody was labelled with ssDNA containing 10 nt unique barcode, UMI (8 nt), stagger sequence (1-16 nt), second UMI (8 nt) and polyA sequences. N^6^-(6-Azido)hexyl-3’-dATP were added to ssDNA oligos (Biolegio) using terminal transferase (TdT, NEB) at 25:1 molar ratio for 2h at 37°C, followed by column purification (Zebaspin RNA clean&concentrator kit) and subsequent coupling to DBCO-PEG12-TCO (Jena Biosciences) at 1:5 molar ratio overnight at room temperature. Following column purification (Zebaspin RNA clean&concentrator kit) to remove excess DBCO-PEG12-TCO, antibody and ssDNA were incubated at a 1:2 molar ratio for 2 hrs at room temperature to allow covalent conjugation. The reaction was quenched by a 300-fold molar excess of 3,6-diphenyl tetrazine. The conjugation efficiency was confirmed by agarose gel electrophoresis. The antibody-DNA conjugate panel was compiled by combining all antibodies at a concentration of 250 ng/ml each in blocking-buffer. The list of antibodies and corresponding ssDNA barcode sequences can be found in Table S1. Extensive validation data for the antibody panel we used is available in ^34,36^.

### RNA and Immuno-Detection

The RAID staining was performed as previously described^33^. Briefly, cultured cells were dissociated, followed by fixation with 2.5 mM DSP and 2.5 mM SPDP in sodium phosphate buffered saline (pH 8.4) for 45 minutes at RT. The fixation was quenched by adding 100 mM Tris-HCl pH 7.5 and 150 mM NaCl for 10 minutes. Subsequently, the cells were blocked and permeabilized for 30 min at RT in a mix of 0.5x PFBB (protein free blocking buffer, Thermo Fisher Scientific) in PBS supplemented with 10 µg/ml tRNA (Roche), 0.5 U/µl RNAsin Plus (Promega) and 0.1% Triton X-100. The panel of ssDNA-labelled antibodies (250 ng/ml for each antibody) was mixed with 2 U/µl RNAsin Plus (Promega) and 0.1% Triton X-100 in PBS:PFBB (1:1) and used to stain the cells. The staining was performed overnight at 4°C on a rotating mixer. To remove unbound antibodies, the cells were washed four times after the staining with a wash-buffer (10% PFBB and 0.05% Triton X-100 in PBS). Subsequently, the cells were washed two more times in 10% PFBB in PBS, followed by single cell sorting (BD FACS Aria) into 384-well plates containing CEL-seq2 primers (Table S10). The library preparation adapted from the previously published protocol^33^. ERCC spike-in (1:100,000 dilution, Thermo Fisher Scientific) was added to the reverse crosslinking mix. The transcriptome and antibody barcodes were separated by performing a double bead-purification after first strand synthesis and pooling. The RNAs were selected using a 0.6x Ampure XP bead ratio (Beckman Coulter) purification, while the antibody barcodes were selected through 2.0x Ampure XP bead ratio purification. The RNAs were subsequently subjected to second strand synthesis and *in vitro* transcription as previously described (Gerlach et al., 2019). The antibody barcode part was first linearly amplified using FWD pre-amplification primer (5’ CACGACGCTCTTCCGATCT 3’) for 10 cycles followed by addition of T7 primer (5’ GCCGGTAATACGACTCACTATAGGG 3’) for 2 cycles to generate dsDNA. The dsDNA generated was then subjected to *in vitro* transcription using Megascript T7 transcription kit (Thermo Fisher Scientific). The amplified RNA from the transcriptome part and antibody barcode part was subjected to reverse transcription primed with random octamer with FWD handle (5’-CACGACGCTCTTCCGATCTNNNNNNNN-3’) and FWD pre-amplification primer (5’-CACGACGCTCTTCCGATCT-3’), respectively. Subsequently, the cDNA was amplified followed by Illumina sequencing index addition as we previously described^33^. The sequencing was performed with Illumina Nextseq 500. The detailed library preparation protocol is provided in the Supplementary Information.

### RAID sequencing data processing, analysis and clustering

The sequence data were demultiplexed using bcl2fastq software (Illumina). RNA sequence data were processed using CELseq2 pipeline ^58^ which includes demultiplexing based on cell barcodes, mapping to GRCh38 using Bowtie2^59^ and UMI counting by HTseq. Antibody-derived tags (ADT) sequence data were subjected to CITE-seq -Count pipeline with the following settings (--max-error 1 --bc_collapsing_dist 1 --umi_collapsing_dist 1). The generated count tables were then analysed as described as follows. Cells in which > 500 and < 6000 genes, < 50% mitochondrial genes and > 20 ADTs were detected, were retained for subsequent analysis to ensure high quality data. The scRNA data were normalised and batch corrected using SCTransform and Seurat integration workflow (Seurat package version 4.0.0)^40^. TMM-based (trimmed mean of M-values, edgeR package ^60,61^ version 3.32.1) normalisation was implemented on the ADT/antibody data, followed by batch correction using ComBat (sva package ^62^ version 3.38.0). Subsequently, the cells were clustered based on nearest neighbours defined using the 3000 most variable genes and visualised using UMAP dimensional reduction following the Seurat version 4 workflow.

### Identification of robust expression programs

There were two strategies implemented to analyse the gene expression based on the obtained scRNA data. First, a cluster-based differential gene expression analysis was performed using the default settings of Seurat version 4 FindAllMarkers with a cut-off of adjusted p-value < 0.001. This generated a list of genes that are differentially expressed in each cluster with regards to the rest of cell populations. The second strategy is a cluster-agnostic approach, which is based on non-negative matrix factorization (NMF). Briefly, log-transformed normalised counts were centred and all negative values were transformed to 0. Afterwards, NMF was performed as previously described in ^41^ using *k* ranging from 5 – 10. The top 50 genes (NMF score-based) of each *k* were selected and defined as expression programs. Furthermore, we selected those with at least 70% overlap with another program detected with different *k*-values. One program with highest overlap was chosen from each set of overlapping programs to avoid redundancies, as previously described in^41^. As the genes obtained from both strategies are quite similar, we combined the output by generating contingency tables using the percentage of overlaps of the gene lists from both strategies. Following clustering, unique genes from the co-clustering programs are selected as robust expression programs. Gene ontology analysis was performed on these lists of genes using MSigDB and Cell Markers options of GeneTrail^63^ (https://genetrail.bioinf.uni-sb.de/) for biological interpretation.

Finally, expression program scores for every cell were calculated. The log-transformed normalised counts were centred and a summation of such counts from each gene in the robust expression program were calculated. To determine the significance of the expression program scores in each of the clusters, control expression program scores were calculated using random cells outside of the cluster of interest, where *n* is the number of cells in the cluster of interest. Significance was calculated using pairwise Wilcoxon rank sum test, between expression program of interest vs control expression program. Alternatively, to determine significance of the expression program scores between experimental conditions, the NT (non-treated) condition is used as control expression program scores in the Wilcoxon rank sum test.

### Antibody-derived tag (ADT) based signalling pathway analysis

A pathway-based approach was used in the analysis of ADT data. ADT were categorised into different pathways based on the biological activity of the antibody target(s). The following pathways are considered: Wnt, BMP, TNFα, TGFβ, Notch, EGF, JAK/STAT, as well as adhesion-related and differentiation-related pathways. The normalised and scaled ADT data were used to calculate a score for each pathway per cell. The scores of molecules that activate a given pathway are summed, whereas those that inhibit the pathway are subtracted from the pathway score. The list of which molecules are used in which pathways are given in Table S7.

### Tolerance score definition and computation

A putative tolerance score was computed based on the ADT data. Briefly, differential expression analysis of the TMM-normalised ADT data was performed using pairwise Kolmogorov-Smirnov test. An ADT marker is defined as a feature with average difference in expression > 0.25 and adjusted p-value < 0.005 in all the possible pairwise comparisons of a given cluster/condition. Based on the observation that SJG26 ADT cluster 2 markers and SJG26 ADT RAG markers are principally similar, these ADT features are defined as “tolerance” markers. The tolerance score was calculated by averaging the normalised and scaled ADT signals of the SJG26 ADT cluster 2 / RAG markers (Table S7).

### Identification of RNA markers of tolerance

Feature selection by random forest followed by validation using stepwise regression based on Akaike Information Criterion (AIC) were employed in identifying genes marking tolerant phenotype. First, highly expressed genes of each patient were selected, in which high expression is defined as top 5%. Highly expressed genes that are present in at least one expression program and also identified as highly expressed in both patients were retained (216 genes). Random forest regression was performed with these 216 genes as independent variables and the tolerance score of both patients as the response variable. The function rfPermute (rf Permute package, version 2.2) was used to run the random forest model with the following parameters; mtry=23, nrep=100, ntree=1000, and importance=TRUE. The validation using stepwise regression was performed using lm (stats package, version 4.0.5) and stepAIC (MASS package, version 7.3-54) functions. The genes obtained in both approaches were then compared. Kaplan-Meier plotter (https://kmplot.com/)^64,65^ were employed to investigate the relation of the identified genes with poor overall survival up to 60 months post-diagnosis.

### RNA Velocity analysis

The raw FASTQ files were subjected to pseudo-alignment to GRCh38 using kb--python (version 0.26.0) ^66,67^. The obtained spliced and unspliced counts were used for RNA velocity analysis using scVelo^42^ Python package (version 0.23.0). The velocity was projected on the RNA and ADT UMAP coordinates generated in Seurat processing of the datasets.

### Inference of CNV from scRNA data

Large copy number variations (CNV) were inferred from the scRNA data using inferCNV package version 1.7.1 (inferCNV of the Trinity CTAT Project. https://github.com/broadinstitute/inferCNV) first described in^38^. scRNA data from non-malignant keratinocytes published in Gerlach et al., 2019 were used as reference in this analysis. The following settings were used in the analysis; cutoff=0.1, HMM=TRUE, analysis_mode=“subclusters”, tumor_subcluster_partition_methods=“random_trees”, denoise=TRUE, BayesMaxPNormal=0.5).

### Bulk RNA sequencing and expressed SNV analysis

Bulk RNA was isolated from the same cell populations subjected to the single cell RAID. The RNA isolation was performed using Quick-RNA Miniprep kit (Zymo Research), following manufacturer’s instructions. Subsequently, sequencing libraries were prepared using KAPA RNA Hyperprep kith with Riboerase (Roche) without alteration to the provided manual. The libraries were sequenced on Illumina Nextseq 500. The obtained bulk RNA sequencing data were demultiplexed with bcl2fastq software (Illumina) and mapped to GRCh38 using STAR version 2.7.9a^68^. The obtained count tables were further analysed using DEseq2 (version 1.30.1)^69^. FASTQ files of bulk RNA sequencing data were mapped to GRCh37 using STAR version 2.7.9a^68^. The genomic coordinates of the SNV of interest were obtained from the list of mutations previously reported in SJG26 and SJG17 patients^35^. Samtools mpileup (version 1.12)^70^ were used to quantify the number of reads mapped as reference or alternative allele on those coordinates. The SNV were further filtered for nonsynonymous SNV that are present in both technical replicates. The read counts of both technical replicates were then merged and filtered further for SNVs with merged counts of > 20 reads. Chi-square tests were employed to determine the significance of reference/alternative allele ratio between selected (AG/R/RAG) and unselected populations (NT).

## Supporting information

RAID protocol

## Data accessibility

The sequencing data has been deposited to the Gene Expression Omnibus (GEO) repository with accession number GSE214145 and GSE214146

## Code availability

Scripts and code used for the single-cell multi-omics analyses are available at https://github.com/dwkarjosukarso/HNSCC_manuscript/.

---

**Supplementary Figure 1:**
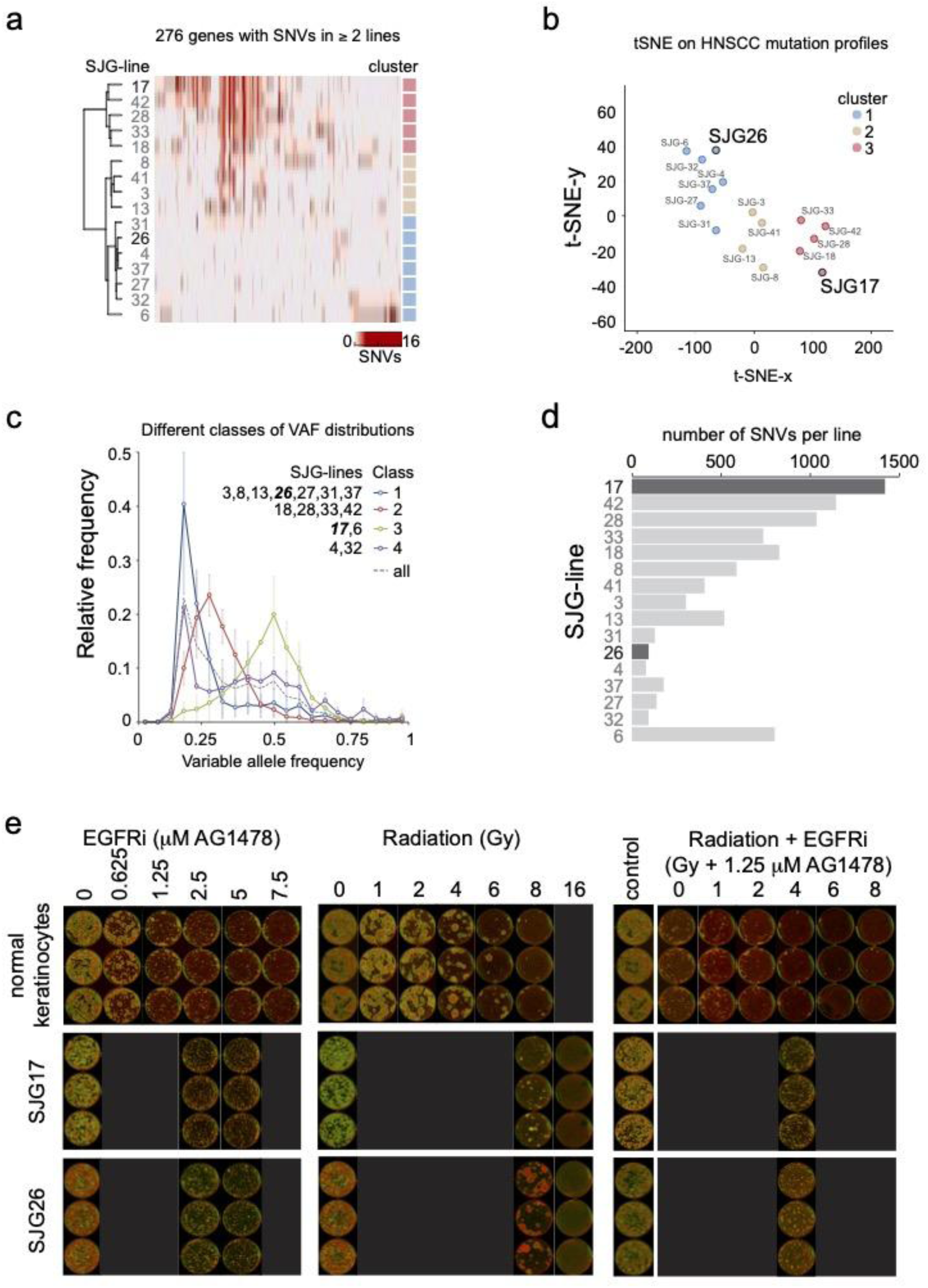
Exome-sequencing data based characterisation and selection of suitable low-passage cultured HNSCC lines. **(a)** Heatmap representing the number of Single Nucleotide Variants (SNVs) called in 276 genes that displayed SNVs in exome sequencing data (REF) from at least two out of the 16 low passage HNSCC cultured cell lines. Data was clustered on both the SJG lines and SNVs. Cluster identity is indicated on the right. **(b)** t-SNE plot of the mutational profiles of the same 276 genes of the collection of 16 SJG lines, colors indicate clusters identified in panel a. Selected SJG26 and SJG17 lines are highlighted. **(c)**Analysis of the distribution of Variable Allele Frequencies (VAFs) for each SGJ line based on exome-sequencing data identifies 4 major classes of VAF distributions. Lines contained in each class are indicated, as are class average profiles, error bars indicate SD for each VAF bin per class. Note that the selected SJG26 and SJG17 lines represent distinct VAF classes. **(d)** Barchart of the absolute number of SNVs per SJG line. Bars representing SJG26 and SJG17 are indicated in dark grey. **(e)** Colony formation assays of normal human epidermal (foreskin) keratinocytes, SJG26 and SJG17 low-passage HNSCC lines treated with a range of concentrations of EGFR inhibitor (AG1478) and/or doses of gamma-radiation (Gy) to identify suitable conditions for selecting tolerant cells from the parental populations.

**Supplementary Figure 2:**
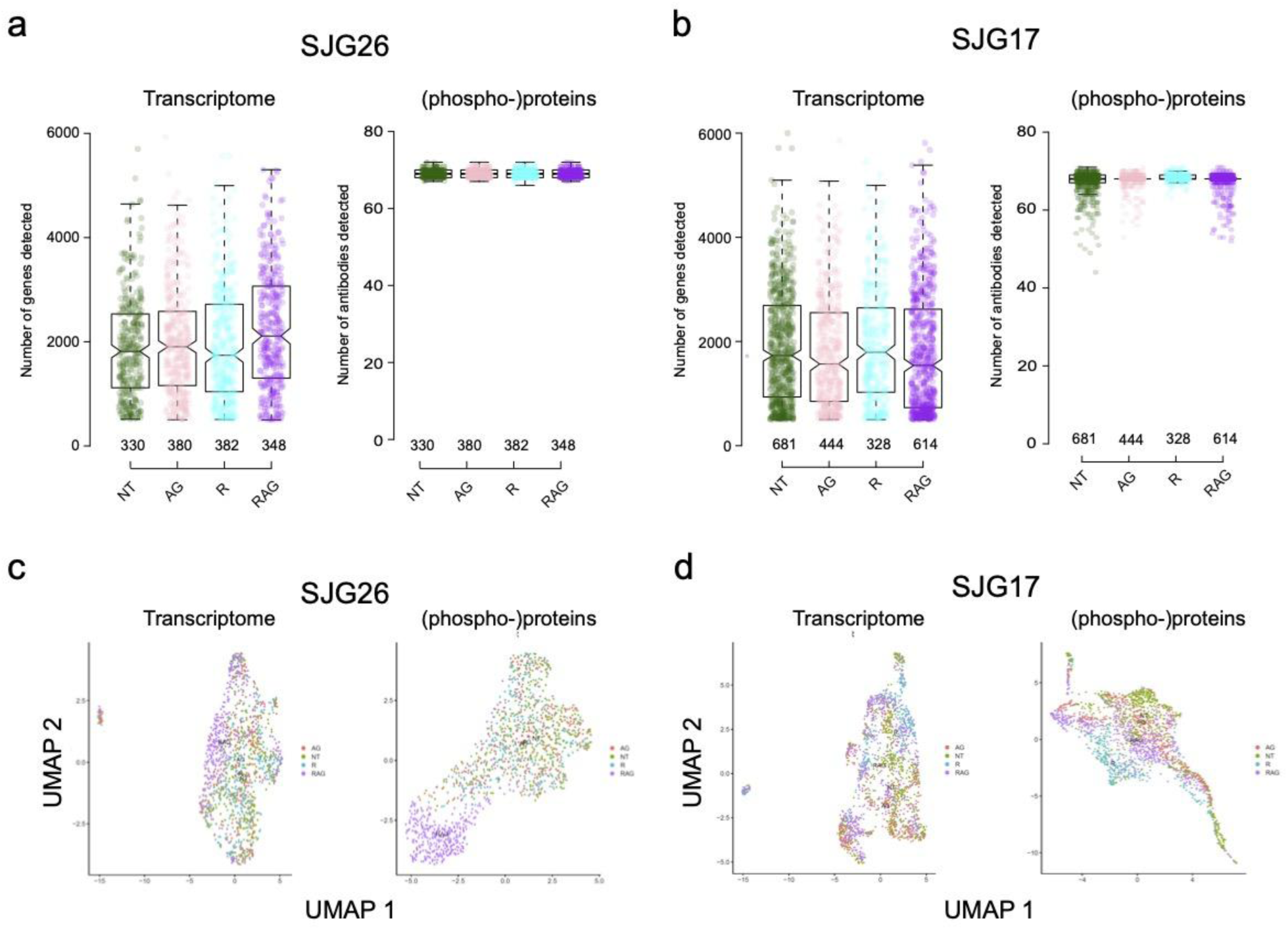
Characteristics of quality and sample distribution of the RNA and Immuno-Detection datasets. **(a,b)** Combined box- and strip plots representing the number of genes and (phospho-)proteins detected from the control and drug-tolerant SJG26 and SJG17 populations, respectively. Each data point represents an individual cell. Boxes indicate 2^nd^ to 3^rd^ quartile, whiskers indicate 1.5x IQR, notches in the box indicate the location of the median. Numbers below the boxes indicate the number of individual cells in the dataset for each population. **(c,d)** UMAP representations of selected drug-tolerant cells from SJG26 or SJG17 HNSCC lines, respectively, based on either RNA transcriptome or antibody-derived (phospho-)protein measurements. Colours indicate the drug-tolerant cell population identities.

**Supplementary Figure 3:**
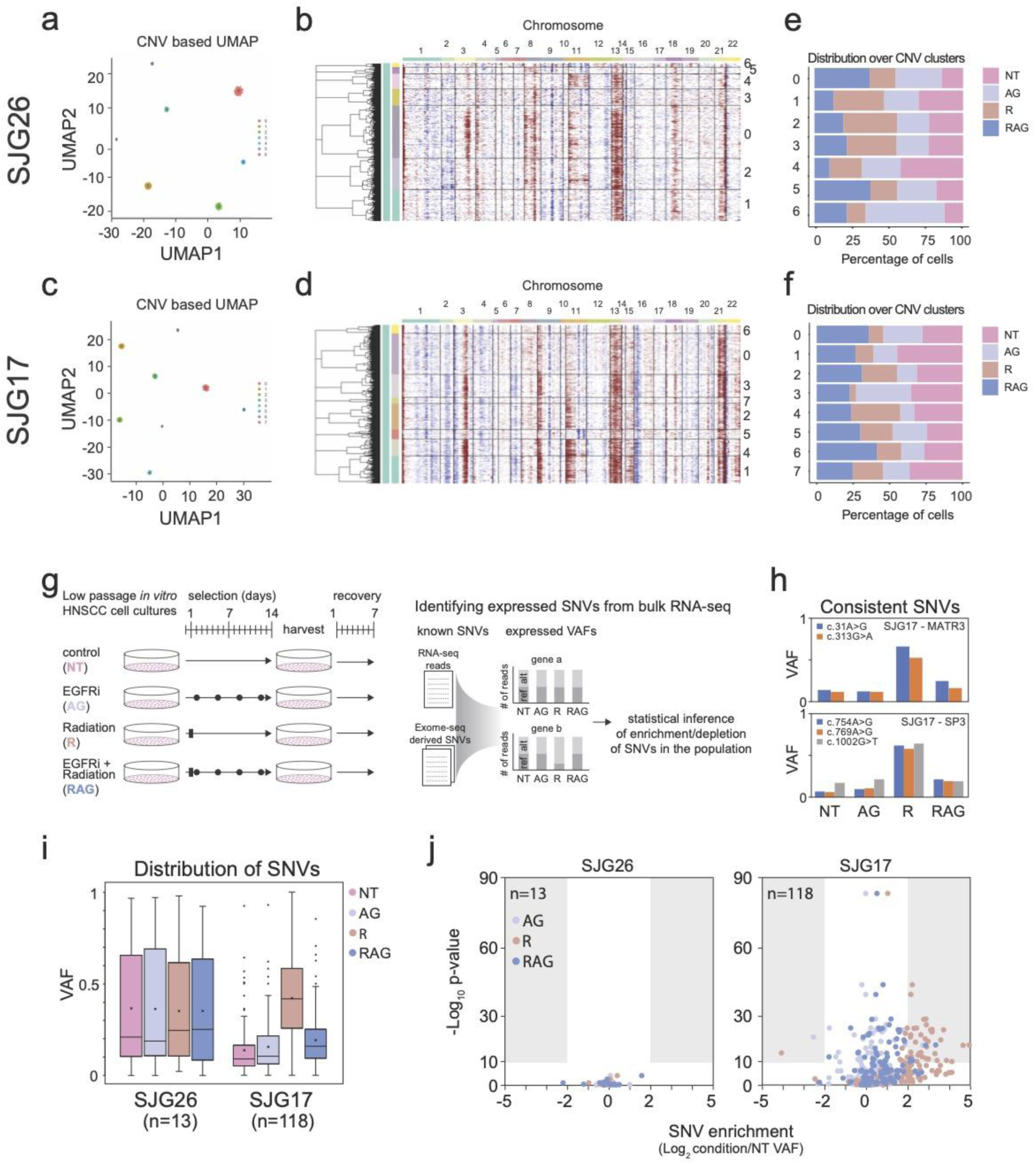
Inferred karyotypic and SNV characterisations of drug tolerant cell populations. **(a)** UMAP representation of inferred Copy Number Variation (CNVs) per cell at low resolution (bin size 100 Mb), identifying 7 distinct clusters of CNV patterns for individual cells from the SJG26 derived drug tolerant populations **(b)** inferred CNV profiles for individual cells from the SJG26 derived drug tolerant populations ordered by genomic location in 1Mb bins and clustered on the cells. **(c,d)** same as **a** and **b**, but for individual cells from the SJG17 derived drug tolerant populations. **(e,f)** Stacked bargraphs of the relative proportions of cells from the different drug tolerant populations for each cluster for SJG26 and SJG17, respectively. **(g)** Schematic representation of the experimental and analysis workflow to assess enrichment of known SNVs from bulk RNA-sequencing data for the selected drug-tolerant populations for SJG26 and SJG17, respectively. **(h)** Bar graph of the VAFs detected in the bulk RNA-seq data distinct SNVs detected in the same gene in the SJG17 drug-tolerant populations. **(i)** Box plots of the VAFs of detected SNVs in the bulk RNA-seq data from the indicated SJG26 and SJG17 populations. N indicates the number of SNVs detected and included in the analysis. Colours indicate the respective populations. **(j)** Volcano plots depicting (log2FC) and −log10 p-value (Chi-squared test) of SNV enrichment in the indicated drug-tolerant populations compared to NT control for SJG26 and SJG17, respectively. N indicates the number of SNVs detected and included in the analysis. Colours indicate the respective populations.

**Supplementary Figure 4:**
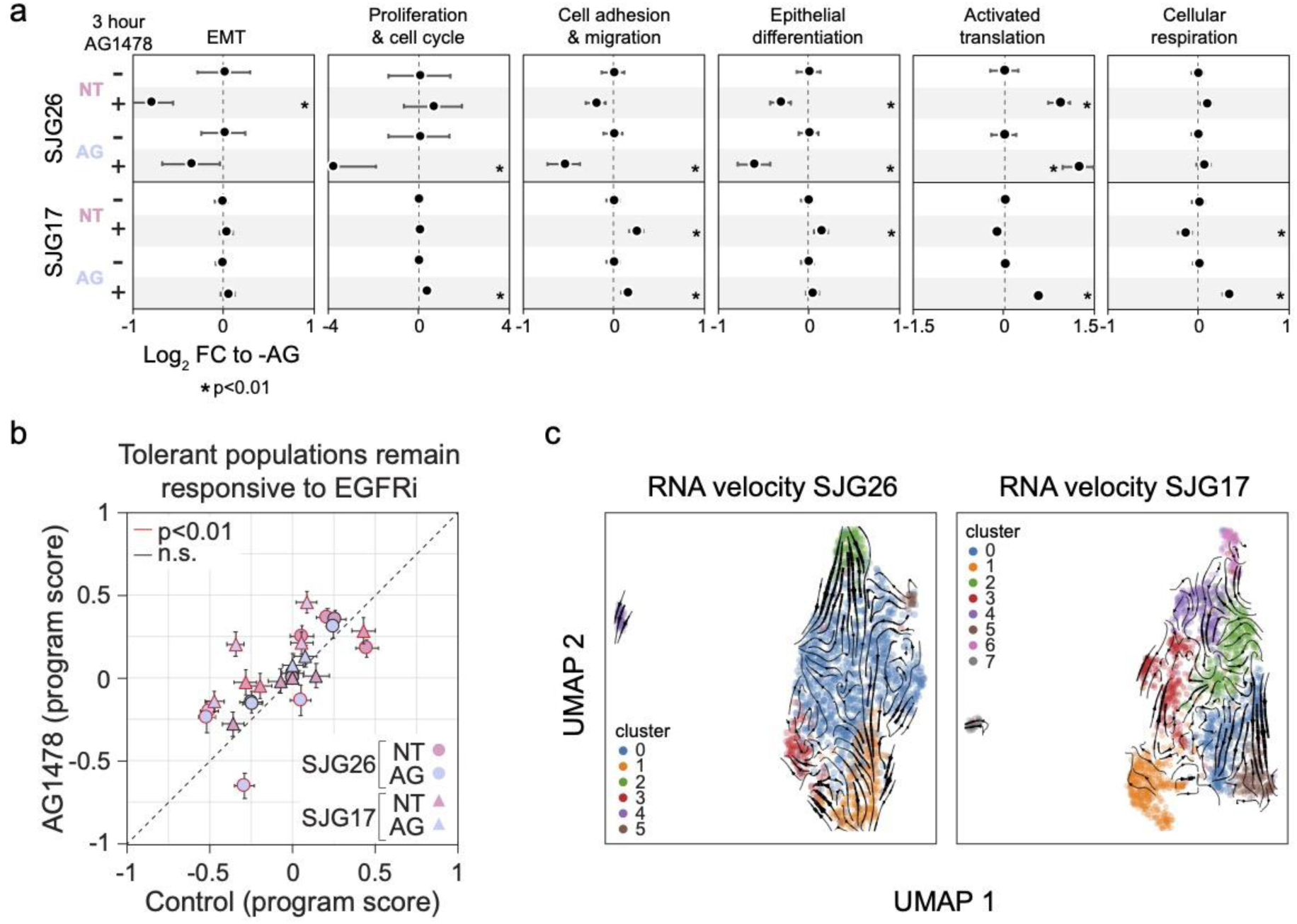
Responsiveness of robust expression programs to EGFRi and RNA velocity analysis **(a)** Most robust expression programs retain some responsiveness to EGFR inhibition in SJG26 and SJG17 HNSCC lines, respectively. Forest plots display the average program score for each indicated population and treatment. Error bars denote 95% CI, * indicates p<0.01, two-sided t-test compared to NT. **(b)** Scatter plot of robust program scores in control and EGFRi conditions for the indicated populations. Symbols with red outline display a statistically significant difference between control and EGFRi conditions (two-sided t-test). Error bars denote 95% CI. **(c)** RNA velocity (scVelo) projected as flow-lines on the RNA derived UMAP for the SJG26 and SJG17 datasets, respectively. Colours indicate clusters called on RNA expression data.

**Supplementary Figure 5:**
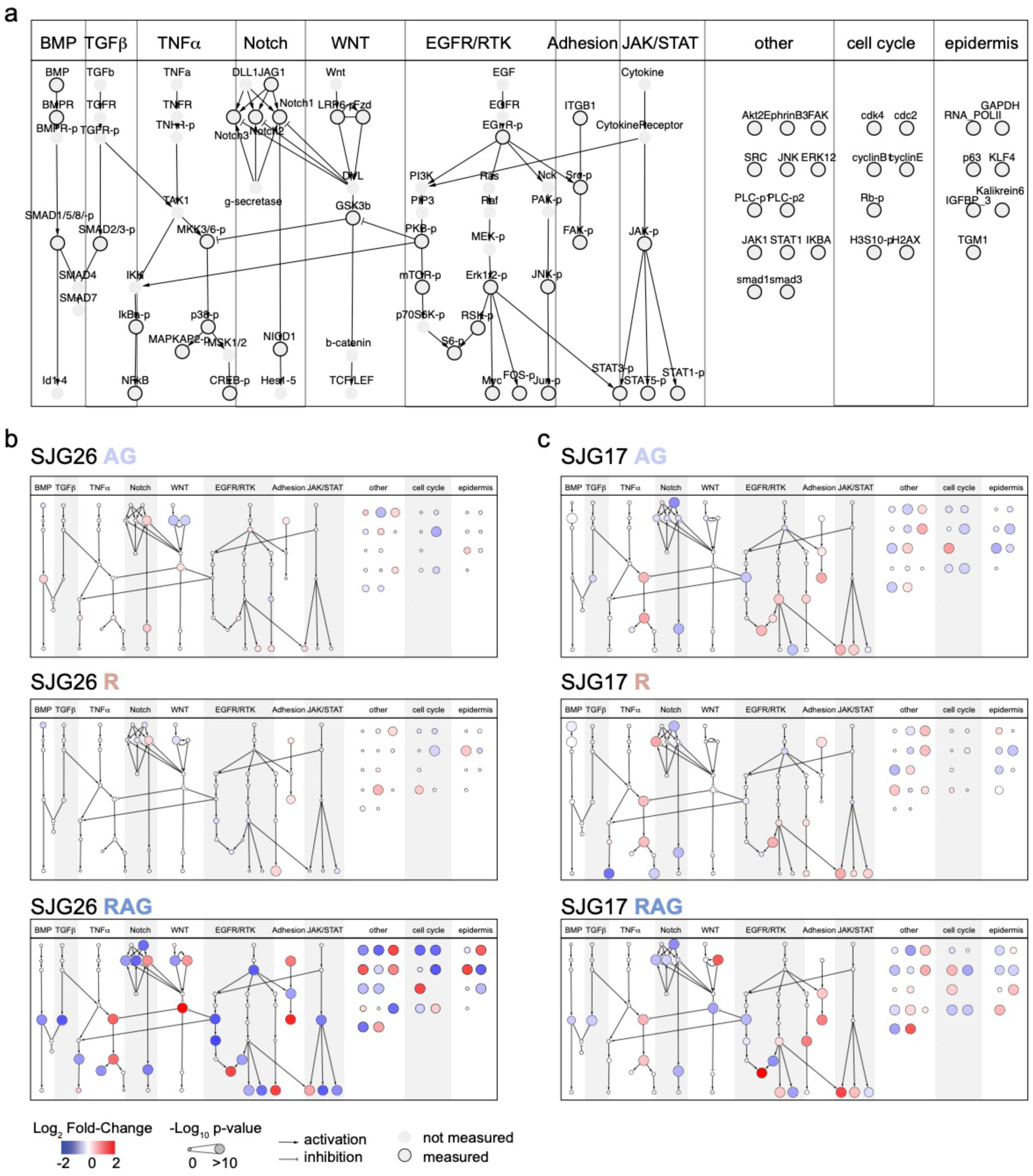
Biochemical network topology and legend. **(a)** Ordered representation of the biochemical network of signaling pathways, including (phospho-)protein names covered by the RAID antibody panel. Arrows depict known kinase-substrate pairs. Colour indicates the effect size and the node size represents its statistical significance (-log_10_ p-value, KS-test). **(b,c)** Differential antibody abundance analyses (control versus indicated population) superimposed the network displayed in **a** for SJG26 **(b)** and SJG17 **(c)**.

**Supplementary Figure 6:**
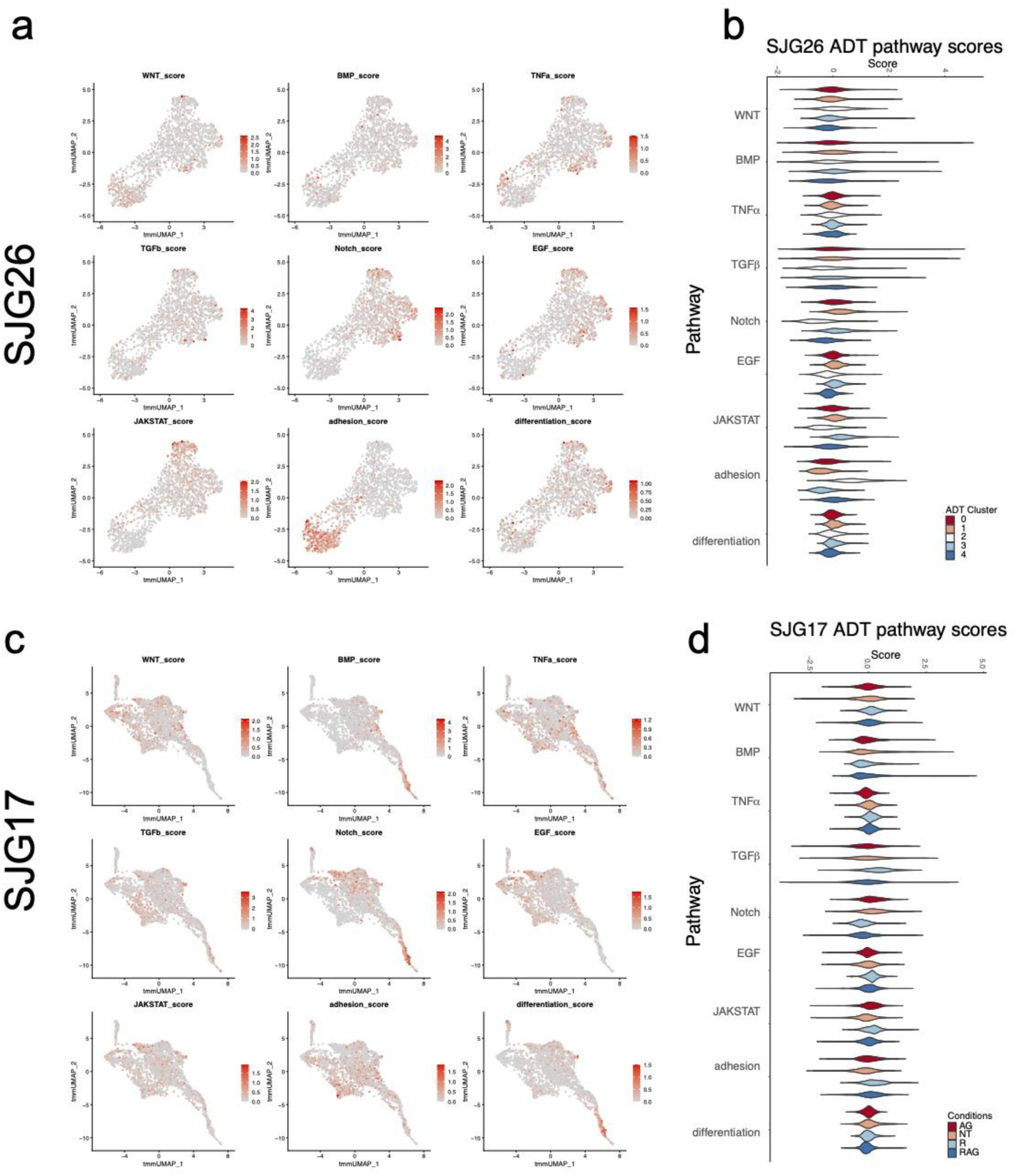
Single-cell antibody-based pathway score analysis. **(a)** Feature plots of indicated pathway scores superimposed on the antibody-data derived UMAPs for the SJG26 dataset. **(b)** Violin plots of pathway scores per (phospho-)protein derived clusters. **(c)** Feature plots of indicated pathway scores superimposed on the (phospho-)protein derived UMAPs for the SJG17 dataset. **(d)** Violin plots of pathway scores per condition.

**Supplementary Figure 7:**
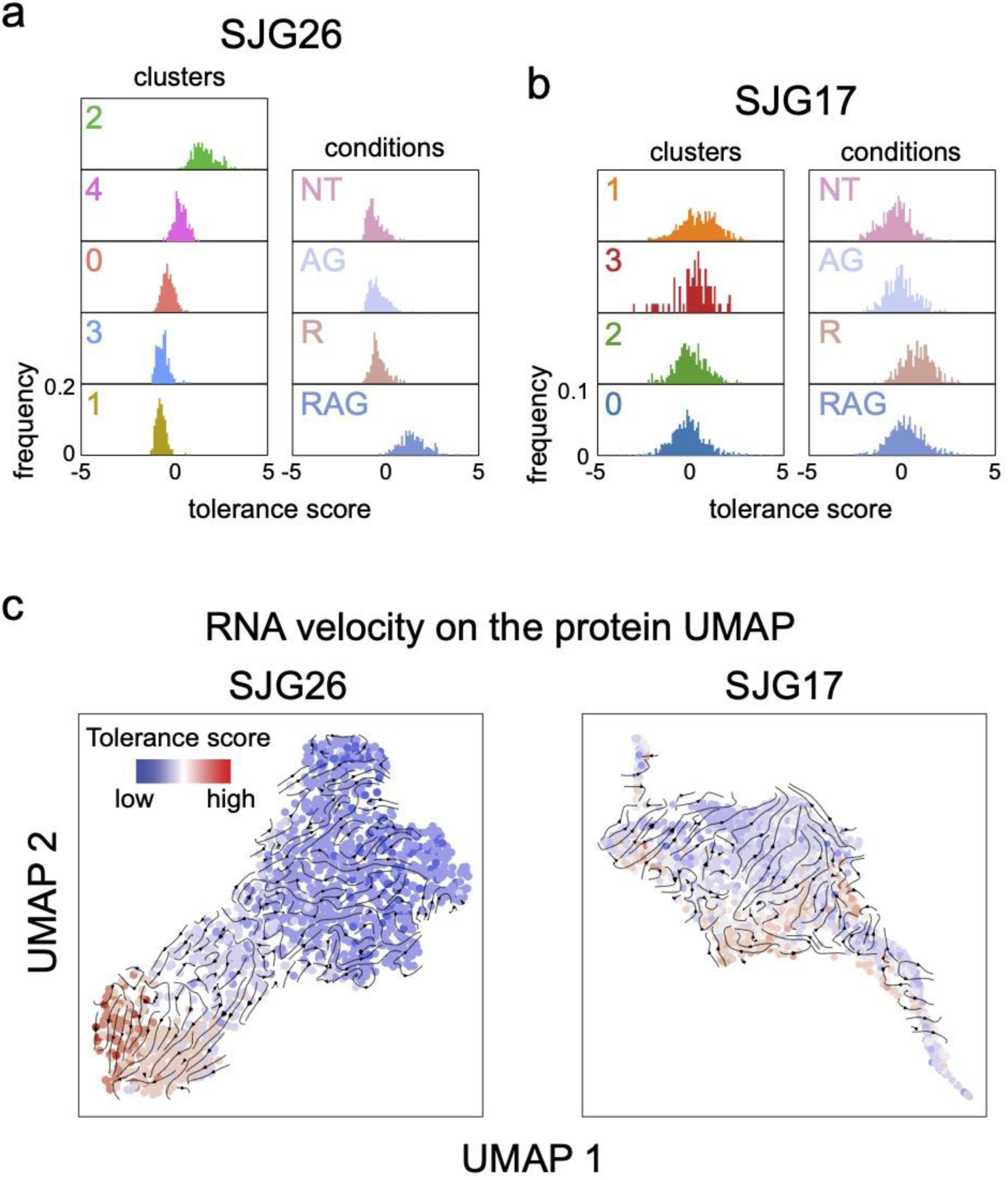
Distribution of the tolerance score over clusters and conditions. **(a)** Histograms of the single-cell tolerance scores in the SJG26 dataset per (phospho-)protein cluster (left panel) and selected drug tolerant populations (right panel). **(b)** Histograms of the single-cell tolerance scores in the SJG17 dataset per (phospho-)protein cluster (left panel) and selected drug tolerant populations (right panel). **(c)** UMAP visualisation of the (phospho-)protein data for the SJG26 and SJG17 datasets with the RNA velocity flowlines indicated as arrows and each cell coloured by their calculated tolerance score.

**Supplementary Figure 8:**
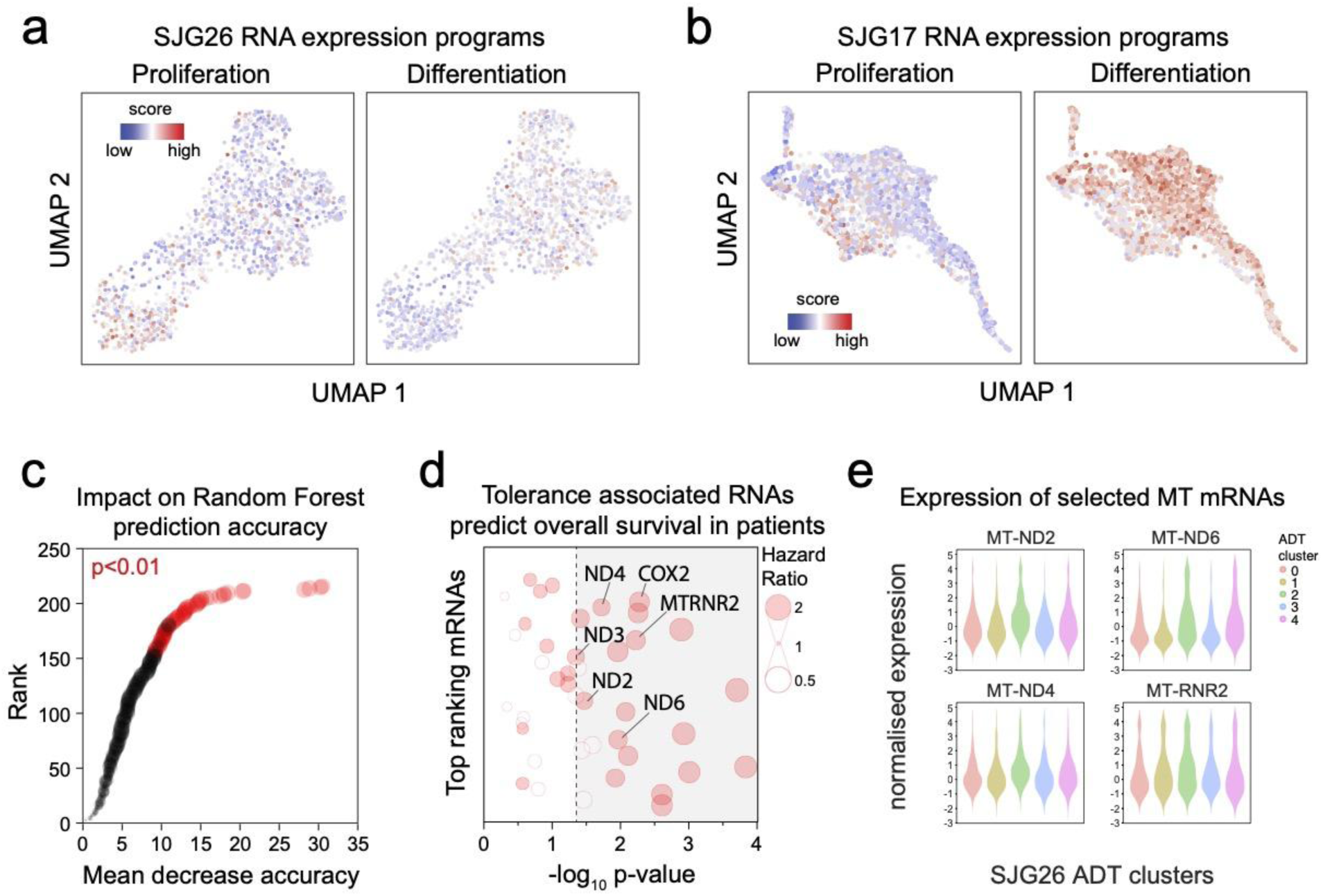
Connecting the tolerance score to transcriptional programs. **(a)** Robust program scores (proliferation, left; differentiation, right) superimposed onto the (phospho-)protein based UMAP for the SJG26 dataset. **(b)** Robust program scores (proliferation, left; differentiation, right) superimposed onto the (phospho-)protein based UMAP for the SJG17 dataset. **(c)**Assessment of the impact of leaving an mRNA out of the Random Forest prediction model. mRNAs ranked by mean decrease accuracy. Colour indicates whether the prediction is significantly affected by the mRNA (empirical p-vale determined with 1000 runs on permuted data). **(d)** Summary of the Hazard ratio and statistical significance of the association of 41 tolerance score associated mRNAs that are also expressed in the TCGA HNSCC cohort dataset. Genes involved in mitochondrial respiration are highlighted. **(e)** Violin plots representing the expression distribution of selected genes involved in mitochondrial respiration in the SJG26 dataset separated by (phospho-)protein based clusters.

**Supplementary Figure 9:**
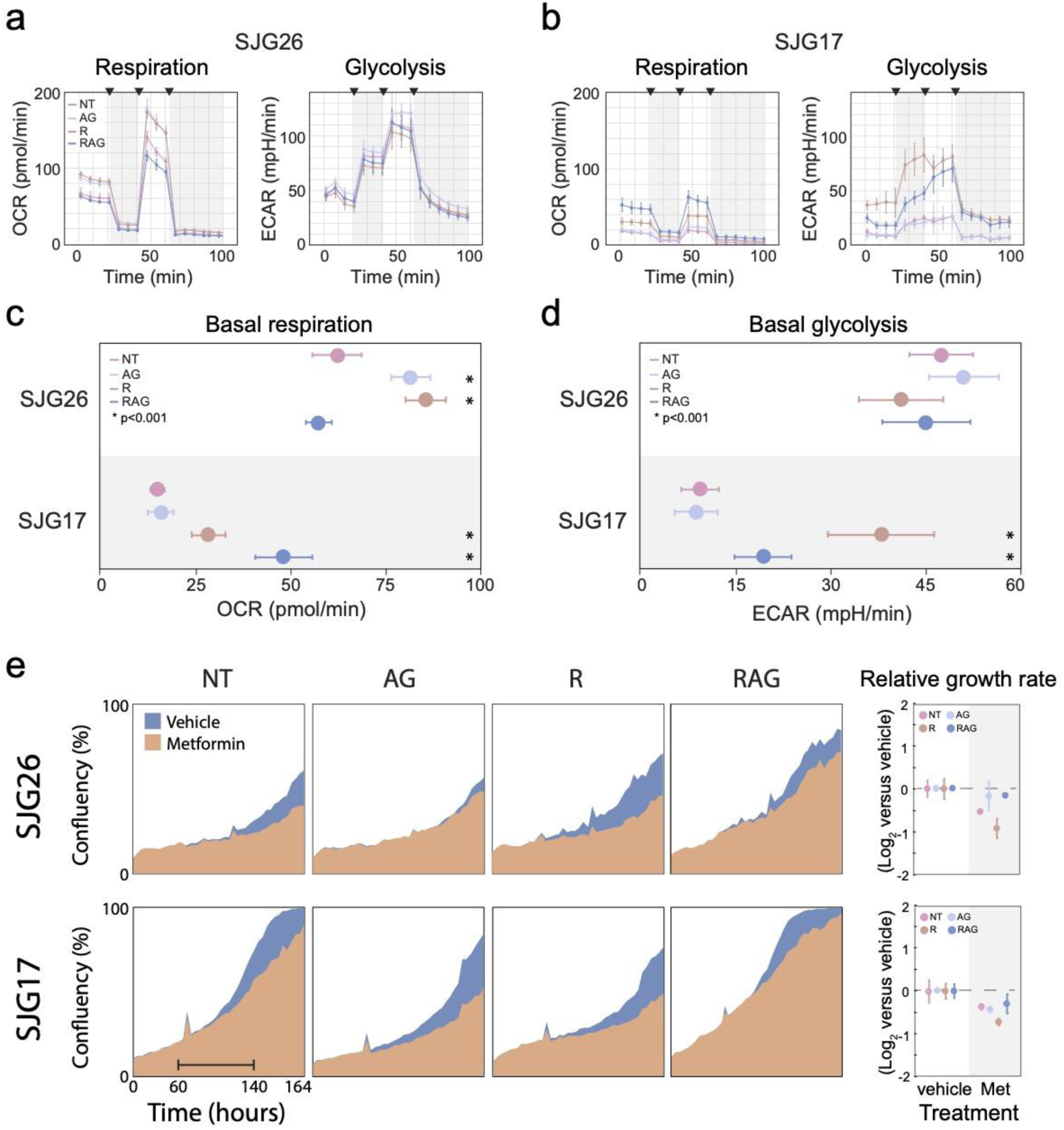
Cellular respiration is associated with drug tolerance. **(a)** Seahorse metabolic profiles of Oxygen Consumption (left) and Extracellular acidification rates (right) to assess respiration and glycolysis, respectively, in the indicated SJG26 populations. N=6 replicates, error bars indicate SD. **(b)** Seahorse metabolic profiles of Oxygen Consumption (left) and Extracellular acidification rates (right) to assess respiration and glycolysis, respectively, in the indicated SJG17 populations. N=6 replicates, error bars indicate SD. **(c)** Basal respiration (OCR) for the indicated populations calculated from the first 4 time-points of the Seahorse analysis in **a**. N=6 replicates per time-point, error bars indicate SD. Asterisk indicates p<0.05, two-sided t-test compared to NT population. **(d)** Basal glycolysis (ECAR) for the indicated populations calculated from the first 4 time-points of the Seahorse analysis in **a**. N=6 replicates per time-point, error bars indicate SD. Asterisk indicates p<0.05, two-sided t-test compared to NT population. **(e)** Live-imaging growth curve analysis (Incucyte) of the indicated populations treated with vehicle or metformin (left panels). Relative growth rate (Log2 transformed) calculated from the exponential growth phase (from 60 to 140 hours as indicated in **a**) for the indicated populations (right panels).

**Supplementary Figure 10:**
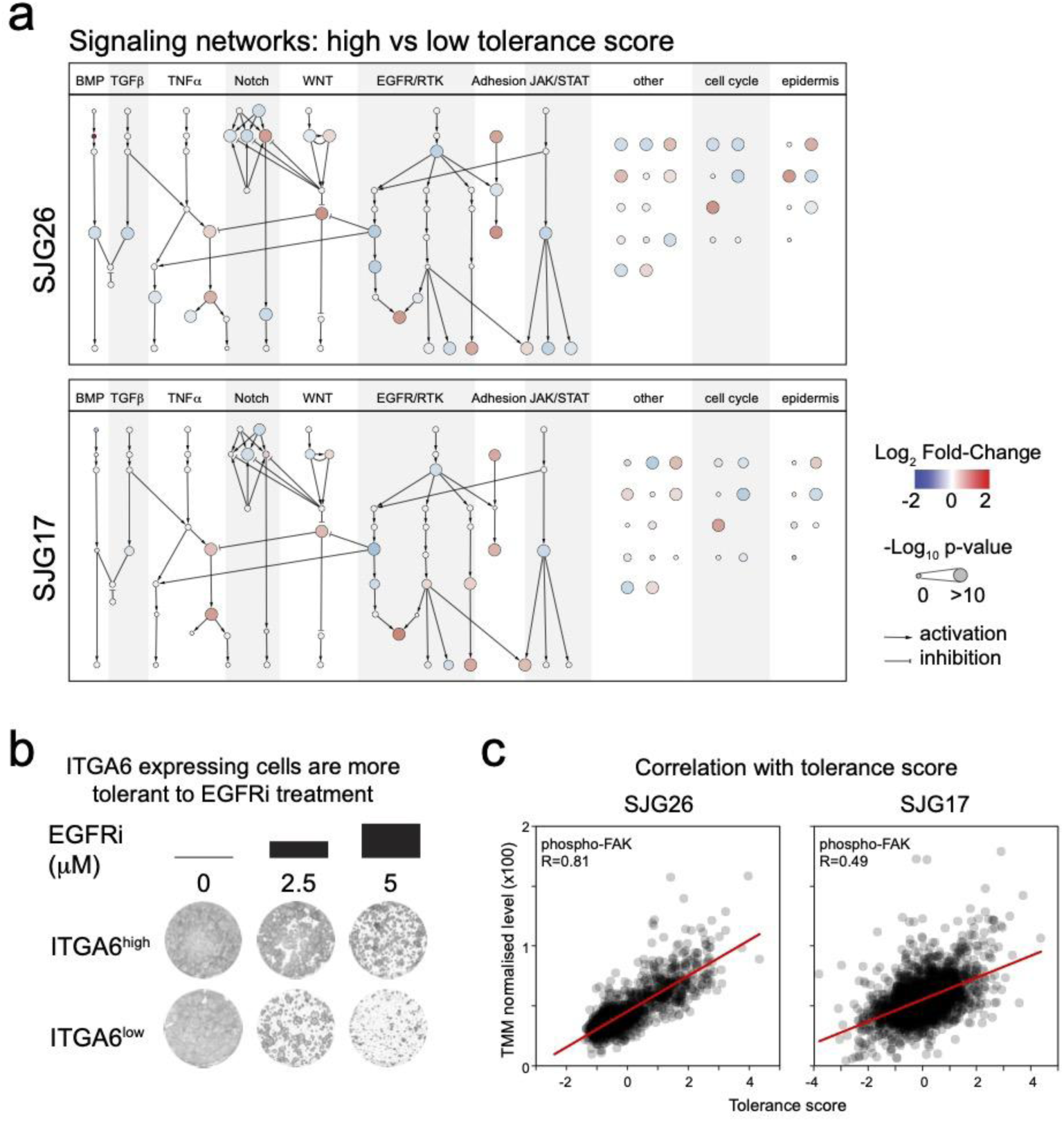
Signaling pathways associated with the calculated tolerance score. **(a)** Differential antibody abundance analyses (cells in the 10^th^ versus 90^th^ percentile of the tolerance score) superimposed the network displayed in Supplementary Figure 5a for SJG26 and SJG17, respectively. **(b)** Representative images of the colony formation assay of cells from the SJG26 parental population prospectively isolated based on their ITGA6 cell surface levels (low versus high) and subjected to different concentrations of EGFRi (AG1478). **(c)** Scatterplots displaying the correlation between phosphorylated FAK (FAK-p) levels and the tolerance score for the SJG26 and SJG17 datasets, respectively. R indicates Pearson’s correlation. Red line represents a linear regression model of the data.

## References

1. Johnson, D. E. et al. Head and neck squamous cell carcinoma. Nat Rev Dis Primers 6, 92 (2020).

2. Ferlay, J. et al. Estimating the global cancer incidence and mortality in 2018: GLOBOCAN sources and methods. Int. J. Cancer 144, 1941–1953 (2019).

3. Bray, F. et al. Global cancer statistics 2018: GLOBOCAN estimates of incidence and mortality worldwide for 36 cancers in 185 countries. CA Cancer J. Clin. 68, 394–424 (2018).

4. Pynnonen, M. A. et al. Clinical Practice Guideline: Evaluation of the Neck Mass in Adults. Otolaryngol. Head Neck Surg. 157, S1–S30 (2017).

5. Picon, H. & Guddati, A. K. Mechanisms of resistance in head and neck cancer. Am. J. Cancer Res. 10, 2742–2751 (2020).

6. Mikubo, M., Inoue, Y., Liu, G. & Tsao, M.-S. Mechanism of Drug Tolerant Persister Cancer Cells: The Landscape and Clinical Implication for Therapy. J. Thorac. Oncol. 16, 1798–1809 (2021).

7. Dhanyamraju, P. K., Schell, T. D., Amin, S. & Robertson, G. P. Drug-Tolerant Persister Cells in Cancer Therapy Resistance. Cancer Res. 82, 2503–2514 (2022).

8. Salem, A. & Salo, T. Identity matters: cancer stem cells and tumour plasticity in head and neck squamous cell carcinoma. Expert Rev. Mol. Med. 25, e8 (2023).

9. Kałafut, J. et al. Shooting at Moving and Hidden Targets-Tumour Cell Plasticity and the Notch Signalling Pathway in Head and Neck Squamous Cell Carcinomas. Cancers 13, (2021).

10. Goyette, M.-A., Lipsyc-Sharf, M. & Polyak, K. Clinical and translational relevance of intratumor heterogeneity. Trends Cancer Res. 9, 726–737 (2023).

11. Van den Bossche, V., et al. Microenvironment-driven intratumoral heterogeneity in head and neck cancers: clinical challenges and opportunities for precision medicine. Drug Resist. Updat. 60, 100806 (2022).

12. Baumeister, P., Zhou, J., Canis, M. & Gires, O. Epithelial-to-Mesenchymal Transition-Derived Heterogeneity in Head and Neck Squamous Cell Carcinomas. Cancers 13, (2021).

13. Meacham, C. E. & Morrison, S. J. Tumour heterogeneity and cancer cell plasticity. Nature 501, 328–337 (2013).

14. Burrell, R. A., McGranahan, N., Bartek, J. & Swanton, C. The causes and consequences of genetic heterogeneity in cancer evolution. Nature 501, 338–345 (2013).

15. Junttila, M. R. & de Sauvage, F. J. Influence of tumour micro-environment heterogeneity on therapeutic response. Nature 501, 346–354 (2013).

16. Brock, A., Chang, H. & Huang, S. Non-genetic heterogeneity--a mutation-independent driving force for the somatic evolution of tumours. Nat. Rev. Genet. 10, 336–342 (2009).

17. Gupta, P. B. et al. Stochastic state transitions give rise to phenotypic equilibrium in populations of cancer cells. Cell 146, 633–644 (2011).

18. Shaffer, S. M. et al. Rare cell variability and drug-induced reprogramming as a mode of cancer drug resistance. Nature 546, 431–435 (2017).

19. Torre, E. A. et al. Genetic screening for single-cell variability modulators driving therapy resistance. Nat. Genet. 53, 76–85 (2021).

20. Emert, B. L. et al. Variability within rare cell states enables multiple paths toward drug resistance. Nat. Biotechnol. (2021) doi:10.1038/s41587-021-00837-3.

21. Goyal, Y. et al. Pre-determined diversity in resistant fates emerges from homogenous cells after anti-cancer drug treatment. bioRxiv 2021.12.08.471833 (2021) doi:10.1101/2021.12.08.471833.

22. Rehmani, H. S. & Issaeva, N. EGFR in head and neck squamous cell carcinoma: exploring possibilities of novel drug combinations. Annals of translational medicine vol. 8 813 (2020).

23. Kalyankrishna, S. & Grandis, J. R. Epidermal growth factor receptor biology in head and neck cancer. J. Clin. Oncol. 24, 2666–2672 (2006).

24. Zhu, X. et al. Prognostic role of epidermal growth factor receptor in head and neck cancer: a meta-analysis. J. Surg. Oncol. 108, 387–397 (2013).

25. Rubin Grandis, J., et al. Levels of TGF-alpha and EGFR protein in head and neck squamous cell carcinoma and patient survival. J. Natl. Cancer Inst. 90, 824–832 (1998).

26. Bonner, J. A. et al. Radiotherapy plus cetuximab for squamous-cell carcinoma of the head and neck. N. Engl. J. Med. 354, 567–578 (2006).

27. Yamaoka, T., Ohba, M. & Ohmori, T. Molecular-Targeted Therapies for Epidermal Growth Factor Receptor and Its Resistance Mechanisms. Int. J. Mol. Sci. 18, (2017).

28. Fasano, M. et al. Head and neck cancer: the role of anti-EGFR agents in the era of immunotherapy. Ther. Adv. Med. Oncol. 13, 1758835920949418 (2021).

29. Puram, S. V. et al. Single-Cell Transcriptomic Analysis of Primary and Metastatic Tumor Ecosystems in Head and Neck Cancer. Cell 171, 1611–1624.e24 (2017).

30. Choi, J.-H. et al. Single-cell transcriptome profiling of the stepwise progression of head and neck cancer. Nat. Commun. 14, 1055 (2023).

31. Yang, Y. et al. Integrated single-cell and bulk RNA sequencing analyses reveal a prognostic signature of cancer-associated fibroblasts in head and neck squamous cell carcinoma. Front. Genet. 13, 1028469 (2022).

32. Kagohara, L. T. et al. Integrated single-cell and bulk gene expression and ATAC-seq reveals heterogeneity and early changes in pathways associated with resistance to cetuximab in HNSCC-sensitive cell lines. Br. J. Cancer 123, 101–113 (2020).

33. Gerlach, J. P. et al. Combined quantification of intracellular (phospho-)proteins and transcriptomics from fixed single cells. Sci. Rep. 9, 1469 (2019).

34. Rivello, F. et al. Single-cell intracellular epitope and transcript detection reveals signal transduction dynamics. Cell Rep Methods 1, 100070 (2021).

35. Hayes, T. F. et al. Integrative genomic and functional analysis of human oral squamous cell carcinoma cell lines reveals synergistic effects of FAT1 and CASP8 inactivation. Cancer Lett. 383, 106–114 (2016).

36. van Buggenum, J. A. G. et al. Immuno-detection by sequencing enables large-scale high-dimensional phenotyping in cells. Nat. Commun. 9, 2384 (2018).

37. van Eijl, R. A. P. M., van Buggenum, J. A. G. L., Tanis, S. E. J., Hendriks, J. & Mulder, K. W. Single-Cell ID-seq Reveals Dynamic BMP Pathway Activation Upstream of the MAF/MAFB-Program in Epidermal Differentiation. iScience 9, 412–422 (2018).

38. Tirosh, I. et al. Dissecting the multicellular ecosystem of metastatic melanoma by single-cell RNA-seq. Science 352, 189–196 (2016).

39. Yang, J., Chen, Y., Luo, H. & Cai, H. The Landscape of Somatic Copy Number Alterations in Head and Neck Squamous Cell Carcinoma. Front. Oncol. 10, 321 (2020).

40. Butler, A., Hoffman, P., Smibert, P., Papalexi, E. & Satija, R. Integrating single-cell transcriptomic data across different conditions, technologies, and species. Nat. Biotechnol. 36, 411–420 (2018).

41. Kinker, G. S. et al. Pan-cancer single-cell RNA-seq identifies recurring programs of cellular heterogeneity. Nat. Genet. 52, 1208–1218 (2020).

42. Bergen, V., Lange, M., Peidli, S., Wolf, F. A. & Theis, F. J. Generalizing RNA velocity to transient cell states through dynamical modeling. Nat. Biotechnol. 38, 1408–1414 (2020).

43. van Buggenum, J. A. G. L. et al. A covalent and cleavable antibody-DNA conjugation strategy for sensitive protein detection via immuno-PCR. Sci. Rep. 6, 22675 (2016).

44. Cancer Genome Atlas Network. Comprehensive genomic characterization of head and neck squamous cell carcinomas. Nature 517, 576–582 (2015).

45. Rêgo, D. F., Pavan, L. M. C., Elias, S. T., De Luca Canto, G. & Guerra, E. N. S. Effects of metformin on head and neck cancer: a systematic review. Oral Oncol. 51, 416–422 (2015).

46. Donati, G. & Watt, F. M. Stem cell heterogeneity and plasticity in epithelia. Cell Stem Cell 16, 465–476 (2015).

47. An, J. S. et al. Integrin alpha 6 as a stemness driver is a novel promising target for HPV (+) head and neck squamous cell carcinoma. Exp. Cell Res. 407, 112815 (2021).

48. Khademi, R. et al. Regulation and Functions of α6-Integrin (CD49f) in Cancer Biology. Cancers 15, (2023).

49. Feng, C. et al. Expression and Prognostic Analyses of ITGA3, ITGA5, and ITGA6 in Head and Neck Squamous Cell Carcinoma. Med. Sci. Monit. 26, e926800 (2020).

50. Zhang, C., Cai, Q. & Ke, J. Poor Prognosis of Oral Squamous Cell Carcinoma Correlates With ITGA6. Int. Dent. J. 73, 178–185 (2023).

51. Dickreuter, E. & Cordes, N. The cancer cell adhesion resistome: mechanisms, targeting and translational approaches. Biol. Chem. 398, 721–735 (2017).

52. Steglich, A., Vehlow, A., Eke, I. & Cordes, N. α integrin targeting for radiosensitization of three-dimensionally grown human head and neck squamous cell carcinoma cells. Cancer Lett. 357, 542–548 (2015).

53. Eke, I., Dickreuter, E. & Cordes, N. Enhanced radiosensitivity of head and neck squamous cell carcinoma cells by β1 integrin inhibition. Radiother. Oncol. 104, 235–242 (2012).

54. Dawson, J. C., Serrels, A., Stupack, D. G., Schlaepfer, D. D. & Frame, M. C. Targeting FAK in anticancer combination therapies. Nat. Rev. Cancer 21, 313–324 (2021).

55. Skinner, H. D. et al. Proteomic Profiling Identifies PTK2/FAK as a Driver of Radioresistance in HPV-negative Head and Neck Cancer. Clin. Cancer Res. 22, 4643–4650 (2016).

56. Pifer, P. M. et al. FAK drives resistance to therapy in HPV-negative head and neck cancer in a p53-dependent manner. Clin. Cancer Res. (2023) doi:10.1158/1078-0432.CCR-23-0964.

57. Goldie, S. J. et al. FRMD4A upregulation in human squamous cell carcinoma promotes tumor growth and metastasis and is associated with poor prognosis. Cancer Res. 72, 3424–3436 (2012).

58. Hashimshony, T. et al. CEL-Seq2: sensitive highly-multiplexed single-cell RNA-Seq. Genome Biol. 17, 77 (2016).

59. Langmead, B. & Salzberg, S. L. Fast gapped-read alignment with Bowtie 2. Nat. Methods 9, 357–359 (2012).

60. Robinson, M. D., McCarthy, D. J. & Smyth, G. K. edgeR: a Bioconductor package for differential expression analysis of digital gene expression data. Bioinformatics 26, 139–140 (2010).

61. McCarthy, D. J., Chen, Y. & Smyth, G. K. Differential expression analysis of multifactor RNA-Seq experiments with respect to biological variation. Nucleic Acids Res. 40, 4288–4297 (2012).

62. Johnson, W. E., Li, C. & Rabinovic, A. Adjusting batch effects in microarray expression data using empirical Bayes methods. Biostatistics 8, 118–127 (2007).

63. Gerstner, N. et al. GeneTrail 3: advanced high-throughput enrichment analysis. Nucleic Acids Res. 48, W515–W520 (2020).

64. Győrffy, B. Discovery and ranking of the most robust prognostic biomarkers in serous ovarian cancer. Geroscience 45, 1889–1898 (2023).

65. Nagy, Á., Munkácsy, G. & Győrffy, B. Pancancer survival analysis of cancer hallmark genes. Sci. Rep. 11, 6047 (2021).

66. Bray, N. L., Pimentel, H., Melsted, P. & Pachter, L. Near-optimal probabilistic RNA-seq quantification. Nat. Biotechnol. 34, 525–527 (2016).

67. Melsted, P. et al. Modular, efficient and constant-memory single-cell RNA-seq preprocessing. Nat. Biotechnol. 39, 813–818 (2021).

68. Dobin, A. et al. STAR: ultrafast universal RNA-seq aligner. Bioinformatics 29, 15–21 (2013).

69. Love, M. I., Huber, W. & Anders, S. Moderated estimation of fold change and dispersion for RNA-seq data with DESeq2. Genome Biol. 15, 550 (2014).

70. Li, H. et al. The Sequence Alignment/Map format and SAMtools. Bioinformatics 25, 2078– 2079 (2009).

