## Supplementary material for "Signalling-state dependent drug-tolerance in head and neck squamous cell carcinoma": RAID protocol

**RAID Library Preparation Protocol (July 2020)**

Materials:
- ERCC Spike-in Mix (Thermo Fisher Scientific, 4456740)
- RNasin Plus (Promega, N2615)
- Maxima H minus Reverse Transcriptase (Thermo Fisher Scientific, EP0753)
- Klenow Fragment (3'→5' exo-) (NEB, M0212L)
- ExoI enzyme (NEB, M0293L)
- Second Strand Buffer (Thermo Fisher Scientific, 10812014)
- E. coli DNA Ligase (NEB, M0205L)
- E. coli DNA Polymerase I (NEB, M0209L)
- RNaseH (Thermo Fisher Scientific, AM2293)
- Herculase II Fusion Polymerase kit (Agilent, 600679)
- MEGAscript T7 Transcription kit (Thermo Fisher Scientific, AMB13345)
- rSAP (NEB, M0371L)
- AmpureXP beads (Beckman Coulter, A63882)

Prepare in advance:
- 10 mM dNTP mix
- 1 M Tris-HCl pH 8
- 1 M DTT, keep frozen
- 10 % Triton X-100
- 5X RT-Buffer (home-made without DTT)
- Beads buffer [20 % PEG8000, 2.5 M NaCl]

CEL-Seq2 primers:
5'GCCGGTAATACGACTCACTATAGGGGTTCAGACGTGTGCTCTTCCGATCTNNNNNNNNCGTCT AATTTTTTTTTTTTTTTTTTTTTTTTTV
* Cell barcode sequence (highlighted) is variable. For a full list of barcodes see Supplementary Table 2.

FWD random octamer:
5’ CACGACGCTCTTCCGATCTNNNNNNNN

FWD pre-amplification primer:
5’ CACGACGCTCTTCCGATCT

REV pre-amplification primer:
5’ GTTCAGACGTGTGCTCTTCCGATC

T7 primer
5’ GCCGGTAATACGACTCACTATAGGG

FWD Library primer: (compatible with Illumina sequencing)
5’ AATGATACGGCGACCACCGAGATCTACACTCTTTCCCTACACGACGCTCTTCCGATCT

REV Library indexing primer: (compatible with Illumina sequencing)
5’
CAAGCAGAAGACGGCATACGAGATATCAGTGTGACTGGAGTTCAGACGTGTGCTCTTCCGATC
* Illumina Index sequence (highlighted) is variable.

Preparation of 384-well plates
- Pipette 5 µl of mineral oil in 384-well plates
- Dispense 100 nl of (7.5 ng/µl) CEL-Seq2 primers in the 384-well plates
- Centrifuge and store at -20 °C until ready for sorting

**Reverse cross-linking and add spike in**

- Thaw the plates
- Add 100 nl of reverse cross-linking mix (prepare fresh) using Nanodrop2 (Bionex) and spin down

| Component | Volume for 1 plate (µl) |
|---|---|
| ERCC-Spike in (1:50k) | 22.2 |
| 10 mM dNTP | 13.5 |
| Nuclease-free water | 0.3 |
| 1 M Tris pH 8 | 3.375 |
| 1 M DTT | 2.025 |
| 10% TX-100 | 0.225 |
| RNasin Plus | 3.375 |

*prepare mix for 1 extra plate to account for dead volume

- Incubate in 384-well PCR machine:
  - 45 minutes at 25 °C
  - 5 minutes at 65 °C
  - continuous 4 °C

**Reverse transcription (RT)**

- Prepare the RT-Enzyme mix as indicated below

| Component | Volume for 1 plate (µl) |
|---|---|
| 5x RT-buffer (home-made, without DTT) | 22.5 |
| 10 mM dNTP | 4.5 |
| Klenow enzyme | 5.625 |
| Maxima H minus RT enzyme | 9 |
| RNasin | 3.375 |

*prepare mix for 1 extra plate to account for dead volume

- Dispense 100 nl in each well using Nanodrop2 (Bionex) and spin down
- Perform the RT reaction:
  - 10 minutes at 25 °C
  - 60 minutes at 50 °C
  - 10 minutes at 85 °C
  - continuous 4 °C

Stopping point – samples can be stored @ -20 °C

**First Strand cDNA pooling and Exonuclease (ExoI) treatment**

- Collect and pool the samples per plate into Eppendorf tube
- Spin down at 13000 rpm for 1 min
- Pipette off and discard as much as possible from the upper (mineral oil) phase without disturbing the lower phase
- Collect the (lower) water phase by piercing through the remaining layer of mineral oil and transfer to new tube

  * It is crucial not to have any mineral oil in the final tube since this can disturb the beads purification.
- Measure the collected aqueous phase and add up to 91 µl with water, to level all samples
- Add 5 µl ExoI and incubate for 20 minutes at 37 °C

**First Strand cDNA cleanup**

- Prewarm Ampure XP Beads to room temperature
- Add 35 µl Ampure XP Beads + 61 µl beads buffer to the samples (1x ratio, for RNA) and transfer to 1.5 ml LoBind Tubes
- Incubate at room temperature for 15 min

- Place on magnetic stand for at least 5 min, until liquid appears clear
- Collect 192 µl supernatant and add 35 µl Ampure XP beads + 61 µl beads buffer (2x ratio for ADT), incubate at RT for 15 min
- Continue with 1x purification beads and wash twice with 200 µl freshly prepared 80 % ethanol
- Air dry beads until completely dry
- Resuspend with 15 µl water. Pipette entire volume up and down to mix thoroughly
- Incubate at room temperature for 5 min
- collect 15 µl in a clean PCR tube
- Do the same with 2x purification beads

Stopping point – Samples can be stored at -20 °C

**FOR RNA PART**

**Second Strand RT reaction**

- Move previous step to ice so it cools below 16 °C
- Add 5.1 µl of the Second Strand mix to each reaction tube

| Component | Volume for 1 sample (µl) |
|---|---|
| Second Strand Buffer | 3.875 |
| 10 mM dNTP | 0.3875 |
| DNA ligase | 0.1395 |
| DNA Polymerase I | 0.5425 |
| RNaseH | 0.1395 |

- Incubate at 16 °C for 120 minutes

**Second Strand cDNA cleanup**

- Prewarm Ampure XP Beads to room temperature
- Add 33 µl (1.6x) Ampure XP Beads to the samples and transfer to 1.5 ml LoBind Tubes
- Incubate at room temperature for 15 min
- Place on magnetic stand for at least 5 min, until liquid appears clear
- Remove and discard the supernatant
- Add 190 µl freshly prepared 80 % ethanol
- Incubate 30 seconds
- Add 190 µl freshly prepared 80 % ethanol
- Incubate 30 seconds
- Air dry beads until completely dry
- Resuspend with 6.5 µl water. Pipette entire volume up and down to mix thoroughly
- Incubate at room temperature for 5 min
- collect 6 µl in a clean PCR tube

Stopping point – Samples can be stored at -20 °C

**FOR ADT PART**

**Library Pre-amplification**

- To the 15 µl purified FS of each sample, add 10 µl of pre-amplification PCR mix:

| Component | Volume for 1 sample (µl) |
|---|---|
| Nuclease-free water | 3 |
| 5x Herculase II reaction buffer | 5 |
| 100 mM dNTP | 0.5 |
| 5 µM FWD pre-amplification primer | 0.5 |
| Herculase II polymerase | 0.5 |

- Amplify using the following PCR conditions:
  - 30 seconds at 95 °C
  - 10 cycles of:
    - 30 seconds at 95 °C

        - 30 seconds at 60 °C
        - 60 seconds at 72 °C
    - 5 minutes at 72 °C
    - Continuous at 4 °C

- Add 0.5 ul of 5 µM T7 primer
- Amplify using the following PCR conditions:
    - 30 seconds at 95 °C
    - 2 cycles of:
        - 30 seconds at 95 °C
        - 30 seconds at 60 °C
        - 60 seconds at 72 °C
    - 5 minutes at 72 °C
    - Continuous at 4 °C

Stopping point – Samples can be stored at -20 °C

**ExoI treatment**

- Add 0.5 µl ExoI and incubate 20 minutes at 37 °C

**Pre-amplifcation cleanup**

- Prewarm Ampure XP beads and beads buffer to room temperature
- Add 26 µl beads + 26 µl beads buffer (2x) to ADT samples
- Incubate at room temperature for 15 min
- Place on magnetic stand for at least 5 min, until liquid appears clear
- Remove and discard the supernatant
- Add 200 µl freshly prepared 80 % ethanol
- Incubate at least 30 seconds, then remove and discard supernatant without disturbing beads
- Repeat this wash step one more time
- Air dry beads until completely dry
- Resuspend with 6.5 µl water
- Incubate at room temperature for 5 min
- Place on magnetic stand for 5 min, until liquid appears clear
- Collect 6 µl in clean PCR tube

**FOR BOTH RNA and ADT PART**

**IVT**

- Prepare the IVT Reaction mix and add 11 µl per sample

| Component | Volume for 1 sample (µl) |
|---|---|
| ATP | 1.7 |
| UTP | 1.7 |
| CTP | 1.7 |
| GTP | 1.7 |
| 10x T7 buffer | 1.7 |
| T7 enzyme | 1.7 |
| RNasin | 0.8 |

- Incubate in a thermal cycler at 37 °C for 14 hours, with lid at 50 °C. Set cycler to go to 4 °C at end of incubation.

**ExoI/rSAP treatment**

- Add 0.5 µl ExoI to the samples
- Add 0.5 µl rSAP to the samples
- Incubate at 25 minutes at 37 °C

**Amplified RNA cleanup**

- Prewarm Ampure XP Beads and beads buffer to room temperature
- Add 32.4 µl (1.8x) to the samples and transfer to room temperature

- Incubate at room temperature for 15 min
- Place on magnetic stand for at least 5 min, until liquid appears clear
- Remove and discard the supernatant
- Add 200 µl freshly prepared 80 % ethanol
- Incubate at least 30 seconds, then remove and discard supernatant without disturbing beads
- Repeat this wash step two more times
- Air dry beads until completely dry
- Resuspend with 12 µl water
- Incubate at room temperature for 10 min
- Place on magnetic stand for 5 min, until liquid appears clear
- Collect 5 µl and store rest at -80 °C

**Library Prep Reverse Transcription**

- To 5 µl amplified RNA add:
  - 1 µl primer
    For RNA part: FWD_8N
    For ADT part: FWD pre-amplification
  - 1 µl 10 mM dNTPs
- Incubate 3 min at 65 °C, quick chill on ice
- Prepare RT-Enzyme mix

| Component | Volume for 1 sample (µl) |
|---|---:|
| 5x First Strand Buffer | 2 |
| RNasin | 0.5 |
| Maxima RT enzyme | 0.5 |

- Add 3 µl of the RT-Enzyme mix to each reaction sample
- Run Reverse Transcription protocol:
  - 10 min at 25 °C.
  - 60 min at 50 °C
  - 10 min at 85 °C
  - continuous at 4 °C

Stopping point – Samples can be stored at -20 °C

**Library Pre-amplification**

- To the 10 µl RT mixture of each sample, add 15 µl of pre-amplification PCR mix

| Component | Volume for 1 sample (µl) |
|---|---:|
| Nuclease-free water | 8 |
| 5x Herculase II reaction buffer | 5 |
| 100 mM dNTP | 0.5 |
| 5 µM FWD pre-amplification primer | 0.5 |
| 5 µM REV pre-amplification primer | 0.5 |
| Herculase II polymerase | 0.5 |

- Amplify using the following PCR conditions:
  - 30 seconds at 95 °C
  - 12 cycles of:
    - 30 seconds at 95 °C
    - 30 seconds at 60 °C
    - 60 seconds at 72 °C
  - 5 minutes at 72 °C
  - Continuous at 4 °C

Stopping point – Samples can be stored at -20 °C

**ExoI treatment**

- Add 0.5 µl ExoI and incubate 20 minutes at 37 °C

**PCR cleanup**

- Prewarm Ampure XP beads and beads buffer to room temperature
- Add 26 µl beads (1x) to the RNA samples and 26 ul beads + 26 ul beads buffer (2x) to the ADT samples
- Incubate at room temperature for 15 min
- Place on magnetic stand for at least 5 min, until liquid appears clear
- Remove and discard the supernatant
- Add 200 µl freshly prepared 80 % ethanol
- Incubate at least 30 seconds, then remove and discard supernatant without disturbing beads
- Repeat this wash step one more time
- Air dry beads until completely dry
- Resuspend with 15 µl water
- Incubate at room temperature for 10 min
- Place on magnetic stand for 5 min, until liquid appears clear
- Collect 15µl in clean PCR tube

**Library Prep. PCR**

- To 15 µl preamplified sample, add 9 µl Library Prep PCR mix and 1 µl of a unique REV Library indexing primer

| Component | Volume for 1 sample (µl) |
|---|---|
| Nuclease-free water | 2.5 |
| 5x Herculase II reaction buffer | 5 |
| 100 mM dNTP | 0.5 |
| 5 µM FWD library primer | 0.5 |
| Herculase II polymerase | 0.5 |

- To each reaction add 1 µl of a unique REV Library indexing primer (2.5 pmol/µl) to index all the samples on the Flow Cell
- Amplify using the following PCR conditions:
  - 30 seconds at 95 °C
  - 6 cycles of:
    - 30 seconds at 95 °C
    - 30 seconds at 60 °C
    - 45 seconds at 72 °C
  - 5 minutes at 72 °C
  - Continuous at 4 °C

Stopping point – Samples can be stored at -20 °C

**ExoI treatment**

- Add 0.5 µl ExoI and incubate 20 minutes at 37 °C

**Purification and size selection**

- Prewarm Ampure XP beads and beads buffer to room temparature
- Add 20.5 µl (0.8x) Ampure XP Beads to the RNA samples and 20.5 µl beads + 29.5 µl beads buffer (2x) to the ADT samples
- Incubate at room temperature for 15 min
- Place on magnetic stand for at least 5 min, until liquid appears clear
- Remove supernatant
- Add 190 µl freshly prepared 80 % ethanol to the beads
- Incubate 30 seconds
- Add 190 µl freshly prepared 80 % ethanol
- Incubate 30 seconds
- Air dry beads until completely dry

- Resuspend in 15 µl water. Pipette entire volume up and down to mix thoroughly
- Incubate at room temperature for 5 min
- Collect samples in a clean LoBind Eppendorf tube

Library prep is now done. The samples should be analyzed by Qbit and Bioanalyzer before sequencing.
